# Aminopeptidase N is a receptor for hedgehog merbecoviruses

**DOI:** 10.1101/2025.09.03.674079

**Authors:** Min Jin, Victoria A. Jefferson, Zhe Zhao, Nicholas J. Catanzaro, Luca De Sabato, Emma L. Keller, Bridget L. Menasche, Cameron Hoffman, Anna Castelli, Ilaria Di Bartolo, Gabriele Vaccari, Tyler N. Starr, Stephanie N. Seifert, Adrian A. Castellanos, Barbara A. Han, Leonid Serebryannyy, Alicen B. Spaulding, Daniel C. Douek, Ana Moreno, Ralph S. Baric, Craig B. Wilen, James M. Rini, Michael Letko

**Affiliations:** Department of Biochemistry, University of Toronto, Toronto, Ontario, Canada; Paul G. Allen School for Global Health, Washington State University, Pullman, WA; Departments of Laboratory Medicine and Immunobiology, Yale University, New Haven, CT; Department of Epidemiology, Gillings School of Global Public Heath, University of North Carolina at Chapel Hill, Chapel Hill, NC; Department of Food Safety, Nutrition and Veterinary Public Health, Istituto Superiore di Sanità, Rome, Italy; Istituto Zooprofilattico Sperimentale della Lombardia e dell’Emilia Romagna (IZSLER) - Virology Department, Via A. Bianchi, 9, 25124 Brescia, Italy; Department of Biochemistry, University of Utah, Salt Lake City, UT; Cary Institute of Ecosystem Studies, Millbrook, NY; Vaccine Research Center, National Institute of Allergy and Infectious Diseases (NIAID), National Institutes of Health (NIH), Bethesda, MD, USA; Department of Molecular Genetics, University of Toronto, Toronto, Ontario, Canada

## Abstract

Merbecoviruses, closely related to the highly pathogenetic Middle East Respiratory Syndrome Coronavirus (MERS-CoV), circulate in hedgehogs throughout Europe and Asia, raising concerns about zoonotic transmission to humans and domestic animals. Unfortunately, how these viruses enter host cells remains unknown, hindering experimental studies. Here, we tested known coronavirus receptor orthologues from European hedgehogs (*Erinaceus europaeus*) and identified Aminopeptidase N (APN) as an entry receptor for hedgehog merbecoviruses. We confirm this result with single-cycle pseudotype and replication-competent virus experiments as well as protein binding assays. A screen of 30 mammalian APN orthologues reveals restricted cross-species receptor use. Cryo-electron microscopy analysis of the viral glycoprotein–receptor complex shows a unique interface distinct from known coronavirus spike:APN interactions, providing a molecular basis for species barriers. These findings expand the known range of receptor use not only within merbecoviruses but also betacoronaviruses, improving our understanding of betacoronavirus receptors, and informing risk assessments for viral emergence.

## INTRODUCTION

Middle East Respiratory Syndrome Coronavirus (MERS-CoV) was first isolated from a pneumonia patient in 2012 and currently has a case fatality rate of 36%^1,2^. Ongoing outbreaks persist due to frequent zoonotic spillover events from dromedary camels to humans, including a recent cluster of cases reported in Riyadh, Saudi Arabia in the spring of 2025^2,3^. Merbecoviruses are a *Betacoronavirus* subgenus that includes the prototypic MERS-CoV and have also been detected in several bat species^4–11^, pangolins^12^, minks^13^, and hedgehogs^14^. Due to the large number of coronaviruses detected in insectivorous bats after the wake of SARS and MERS emergence, Corman et al. hypothesized in 2014 that other insectivorous mammals, particularly hedgehogs, could harbor coronaviruses in a similar manner^14–17^. Since the initial identification of merbecoviruses from European hedgehogs, *Erinaceus europaeus*, in Germany^14^, ErinCoVs have been further identified in France^18,19^, the United Kingdom^20^, China^21,22^, Italy^23,24^, Poland^25^, Russia^26^, and Portugal^27^. Hedgehogs are a likely reservoir for merbecoviruses due to the high rates of detection of viral genomic material in hedgehog sample populations, ranging from 10% in the United Kingdom, to around 60% in Germany and Italy^14,20,23^. Given the high prevalence of merbecoviruses in hedgehogs and the density of these animals across populated urban areas, there are concerns that these animals pose an infectious disease risk to humans, especially as food availability and reduced predation increase the density of hedgehog populations in urban areas^28–30^. Unfortunately, much about ErinCoV virology, including the host receptor(s) employed by these viruses for cell entry, is completely unknown, preventing efforts to study ErinCoVs under experimental conditions. Thus, the risk posed by ErinCoVs to human populations is speculative but uncertain.

Receptor binding has been shown to be an important species barrier for coronaviruses, as many animal species that are otherwise refractory to infection, can be rendered susceptible to coronaviruses if provided with the appropriate host cell receptor^31–34^. To infect host cells, coronaviruses use the viral spike glycoprotein that binds the host cell receptor via its receptor binding domain (RBD)^35,36^. Following receptor binding, the viral spike protein is cleaved by host proteases, and subsequent conformational changes provide the driving force for membrane fusion and viral entry^37–39^. Very few host protein receptors have been reported for the coronaviruses, with multiple viruses using the same receptors^36^. For example, the alphacoronavirus HCoV-NL63 uses human Angiotensin Converting Enzyme 2 (ACE2) as do the betacoronaviruses SARS-CoV and SARS-CoV-2^40–42^, while the host molecule, aminopeptidase N (APN) is widely used by many of the alphacoronaviruses including HCoV-229E^43^. MERS-CoV was initially unique in its reported use of dipeptidyl peptidase IV (DPP4). However, the identification of other DPP4-using merbecoviruses quickly expanded our understanding of the use of this receptor^44,45^. Embecoviruses are a *Betacoronavirus* subgenus that can use glycoconjugates containing 9-*O*-acetylated *N*-acetylneuraminic acid as receptors, and they can use protein receptors, namely, transmembrane serine protease 2 (TMPRSS2) for HCoV-HKU1^46^, carcinoembryonic antigen-related cell adhesion molecule 1 (CEACAM1) for MHV^47^, and dipeptidase 1 (DPEP1) for PHEV^48^. Embecoviruses also possess hemagglutinin esterase (HE), a viral membrane protein and receptor destroying enzyme which may also influence tropism^49–51^. For coronaviruses without an HE gene, only ACE2, DPP4, and APN have been reported as receptors to date, although the receptors of many coronavirus lineages remain undetermined.

In a previous study we showed that the receptor binding for the MERS-CoV spike could be modified by exchanging the RBD from other merbecoviruses^52^. Using this tool, we established that merbecoviruses can be divided into 4 separate functional clades based on indels, sequence features, and phylogenetic similarity^52^. Merbecoviruses with clade 1 RBDs include viruses such as MERS-CoV and HKU4 and have been shown to use DPP4 from diverse host species^12,45,53–55^. Clade 2 merbecoviruses are found in bats across Europe and Asia, comprise the bulk of published wildlife derived merbecovirus sequences, and have been shown to use ACE2 from their respective host species but have limited cross-species receptor recognition^52^. Clade 3 merbecoviruses have been reported in African bat species and can utilize ACE2 from a broad species range including humans^56^. Merbecoviruses with clade 4 RBDs have been identified in both European and Asian hedgehog species and have no ascribed receptor^14,27,52^.

Here we screened potential host receptors for their ability to mediate clade 4 RBD merbecovirus entry. Using our MerbecoType pseudotype platform, we show that aminopeptidase N (APN), a common receptor for alphacoronaviruses, is capable of mediating cell entry for spike genes from hedgehog merbecoviruses that have been found across Europe, including viruses reported in the UK, Germany, Italy, Poland, and Russia. Notably, we show this viral-host interaction is highly species restricted, with APN from very few species capable of supporting viral entry. We further demonstrate a direct interaction between ErinCoV RBDs and hedgehog APN through recombinant protein binding assays and using cryo-EM provide a structural rationalization for the interaction and limited species receptor usage. The structures of complexes between hedgehog APN and the RBDs from both German and Italian ErinCoVs show that the ErinCoV RBD:APN interaction is unique and distinct from that of other coronavirus interactions with APN^57–60^. Structure guided mutagenesis identified regions of the interface which are less tolerant to mutations than other regions, providing further evidence of a species barrier at the level receptor binding for ErinCoVs. Alongside DPP4 and ACE2 which have previously been shown to be merbecovirus receptors, these results uncover a third receptor for the merbecoviruses unique to hedgehogs and the first reported instance of APN use among betacoronaviruses. Collectively, the findings presented here broaden our understanding of coronavirus receptor use and will be instrumental in risk assessment of novel coronavirus discoveries in wildlife.

## RESULTS

### Diverse ErinCoV RBDs infect cells expressing hedgehog APN

We previously screened merbecovirus RBDs for cell receptor binding, and this included testing ErinCoV RBDs with ACE2 and DPP4 orthologues from hedgehogs^34,52^. Although we did not observe clear viral entry with the ErinCoV RBDs, we did detect a faint but reproducible entry signal in some cell lines^52^. Following up on these results, we produced a small panel of viral pseudotypes containing chimeric MERS-CoV spikes with the RBDs from ErinCoVs that produced a non-specific entry signal in our previous study (ErinCoV-12-19, ErinCoV-1-18) as well as those that showed no entry at all (ErinCoV-19-18). Because APN is the only other protein receptor that has been reported for coronaviruses without an HE gene, we wondered if this molecule may serve as the missing receptor for clade 4 merbecovirus RBDs and utilized European hedgehog APN for our following screens. As a control, we also included an alphacoronavirus, HCoV-229E, in our panel because this virus uses human APN as its receptor (**Figure 1A**). To our surprise, several of the ErinCoV RBDs mediated infection in cells expressing hedgehog APN. This outcome appeared to be species specific, as HCoV-229E could only infect cells expressing human APN and ErinCoV RBDs could only infect cells expressing hedgehog APN (**Figure 1B**). To confirm the phenotype, we changed the target cells from 293T to BHK, as BHK cells are not susceptible to coronavirus entry other than HCoV-OC43^61^, and exhibit no detectable background entry signal in our assays^52^. We also re-tested additional human and hedgehog receptor candidates but only observed ErinCoV RBD entry in cells that were transfected with hedgehog APN (**Figure 1C**). The addition of trypsin during the infection amplified the entry signal but did not alter the receptor usage, as the entry signal was still receptor-dependent (**Figure 1D**).

**Figure 1.**
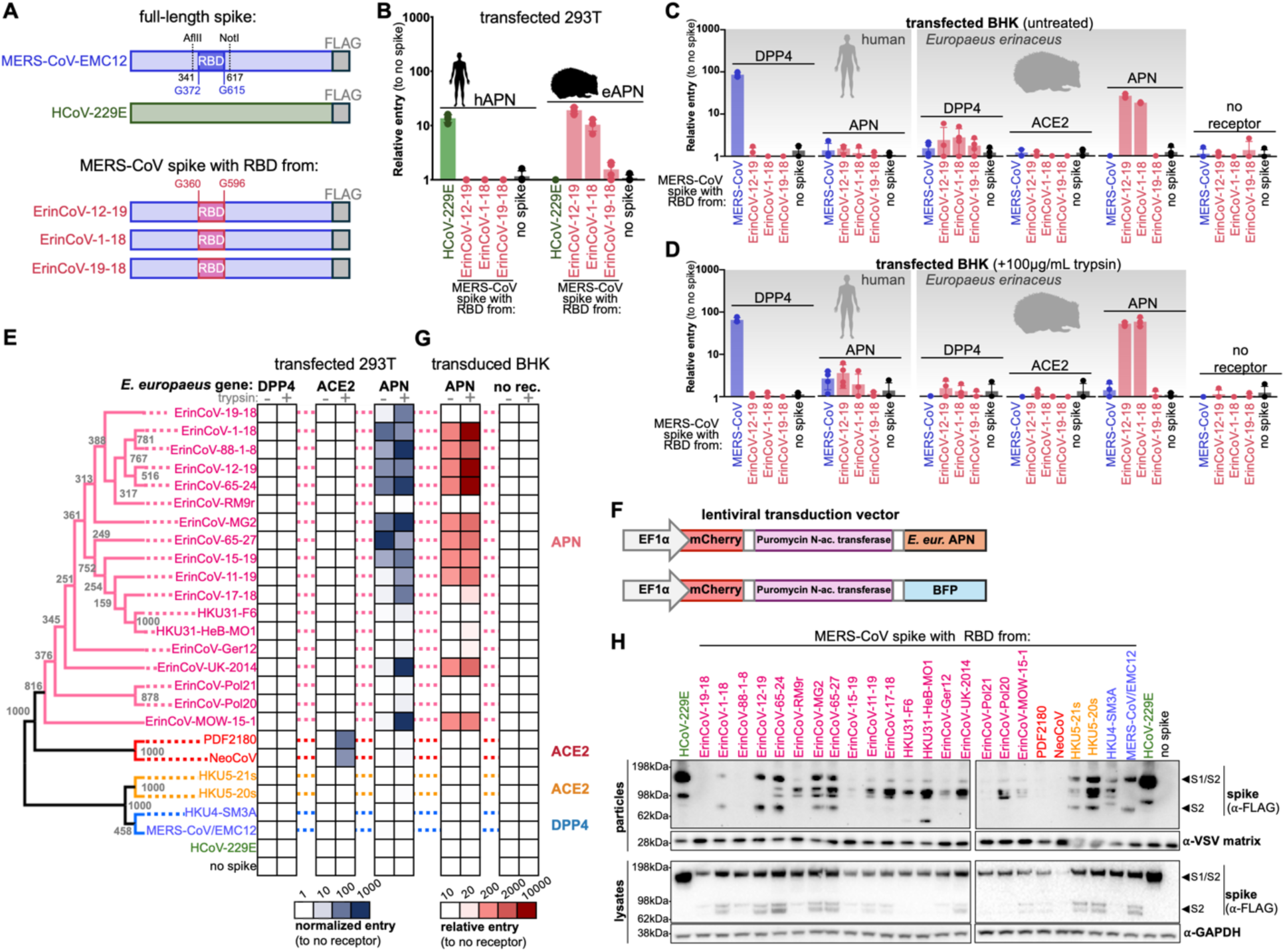
APN from hedgehogs mediates cells entry of diverse ErinCoV RBDs. **(A)** Schematic overview of chimeric and wild-type spikes used in these assays **(B)** 293T cells were transfected with indicated receptors and infected with pseudotypes bearing chimeric spikes with ErinCoV RBDs or HCoV-229E spike and luciferase was measured for cell entry. Infections were performed in four technical replicates. **(C)** BHK cells were transfected with indicated receptors and infected with pseudotypes bearing chimeric spikes with ErinCoV RBDs or MERS-CoV spike and luciferase was measured for cell entry. **(D)** Trypsin was supplemented during the infection. Infections were performed in four technical replicates. **(E)** 293T cells transfected with indicated receptors were infected with pseudotypes bearing chimeric MERS-CoV spike pseudotypes in quadruplicate. **(F)** Schematic overview of lentiviral transduction vector used to produce cells that stably express hedgehog APN **(G)** BHK cells transduced to stably express hedgehog APN were infected with pseudotype bearing chimeric MERS-CoV spike pseudotypes **(H)** Western blot for spike and loading control proteins of lysates and concentrated supernatants from cells producing indicated chimeric MERS-CoV spike chimera pseudotypes.

To assess which hedgehog merbecoviruses use APN as a receptor, we generated a large merbecovirus RBD chimeric spike pseudotype panel, which included all the published variations of the ErinCoV RBDs as well as two representative RBDs from each of the other merbecovirus clades (**Figure 1E; Supplemental Figure 1**). As described above, the ErinCoV RBDs were only capable of infecting 293T cells expressing hedgehog APN but not hedgehog DPP4 or ACE2. Notably, the majority of ErinCoVs tested showed entry with APN, including the viruses from Italy (e.g., ErinCoV-12-19), the UK (ErinCoV-UK-2014), and Russia (ErinCoV-MOW-15-1) (**Figure 1E**). To further confirm these results, we generated a lentiviral vector to transduce BHK cells with hedgehog APN and repeated the entry assay (**Figure 1F, G**). Western-blotting for the FLAG tag on the chimeric spikes showed similar expression levels in cells producing viral pseudotypes (**Figure 1H**). While varying levels of spike were detected in concentrated viral supernatants, these levels did not correlate with viral entry, as previously shown (**Figure 1H**).

The body temperature in hedgehogs is lower than humans or bats, with an average of 35°C^62^. Because previous work has shown that coronaviruses adapt to their environmental temperatures, we tested whether we could further optimize our assays by modulating the pseudotype production temperature. As a control, we included the spike gene from HCoV-229E, which has been shown to yield more infectious pseudotypes when produced at 34°C^63^. In agreement with this earlier work, HCoV-229E spike pseudotypes were more infectious in human APN expressing cells, when produced at 34°C than 37°C. Similarly, ErinCoV RBD chimeras and wild-type spike pseudotypes were more infectious in hedgehog APN cells produced at lower temperatures (**Supplemental Figure 2**).

### Validation of APN-mediated entry with full-length spikes in stably transduced cells

We have shown in previous studies that pseudotyped viruses containing chimeric spikes faithfully reproduce the entry phenotypes of pseudoviruses with wild-type spikes, viruses including replication competent molecular clones, and wild-type viral isolates from field samples^52,64–66^. To further validate the APN-dependent entry we observed with ErinCoV RBD chimeric spikes, we generated wild-type spike proteins from two viruses that produced clear entry signals with APN (ErinCoV-12-19, ErinCoV-UK-2014) and one from a virus that did not (ErinCoV-Ger12 from Germany) (**Figure 1E, G; Figure 2A**). Both chimeric and wild-type spikes expressed similarly in pseudotype producer cells suggesting similar stability levels, however as noted in our previous experiments, the detected levels of spike incorporation varied but did not correlate with entry phenotypes (**Figure 2B**). Pseudotyped particles bearing chimeric spikes and wild-type spikes readily infected both 293T and BHK cells that were transduced to stably express hedgehog APN but not cells that lacked this receptor (**Figure 2C, D**). As we have noted for sarbecoviruses and other merbecoviruses in our previous studies, the chimeric ErinCoV spikes generally entered cells with higher efficiency than the wild-type ErinCoV spikes, which we attribute to the underlying compatibility of the MERS-CoV spike backbone with the cell culture models used in these assays (**Figure 2C, D**)^52,64,67^. It Is also possible that the RBDs in the chimeric spikes have a greater tendency to access the “up” conformation which would facilitate receptor interactions^52,68,69^. To further validate our entry results, we cloned the spike gene from ErinCoV-12-19 into a replication competent Vesicular Stomatitis Virus (VSV)-based fluorescent reporter system (**Figure 2E**)^70^. In agreement with our chimeric and wild-type spike results, cells transduced with hedgehog APN supported clear replication of the ErinCoV-12-19 pseudotyped VSV, with strong GFP and luciferase activity accumulating over the experiment (**Figure 2F, G**). Notably, antibodies directed toward human APN were unable to cross-react with hedgehog APN (**Figure 2H**). Taken together, these results demonstrate that cell lines from different species (humans and hamsters) can be rendered susceptible to cell entry by ErinCoV spikes when provided with hedgehog APN.

**Figure 2.**
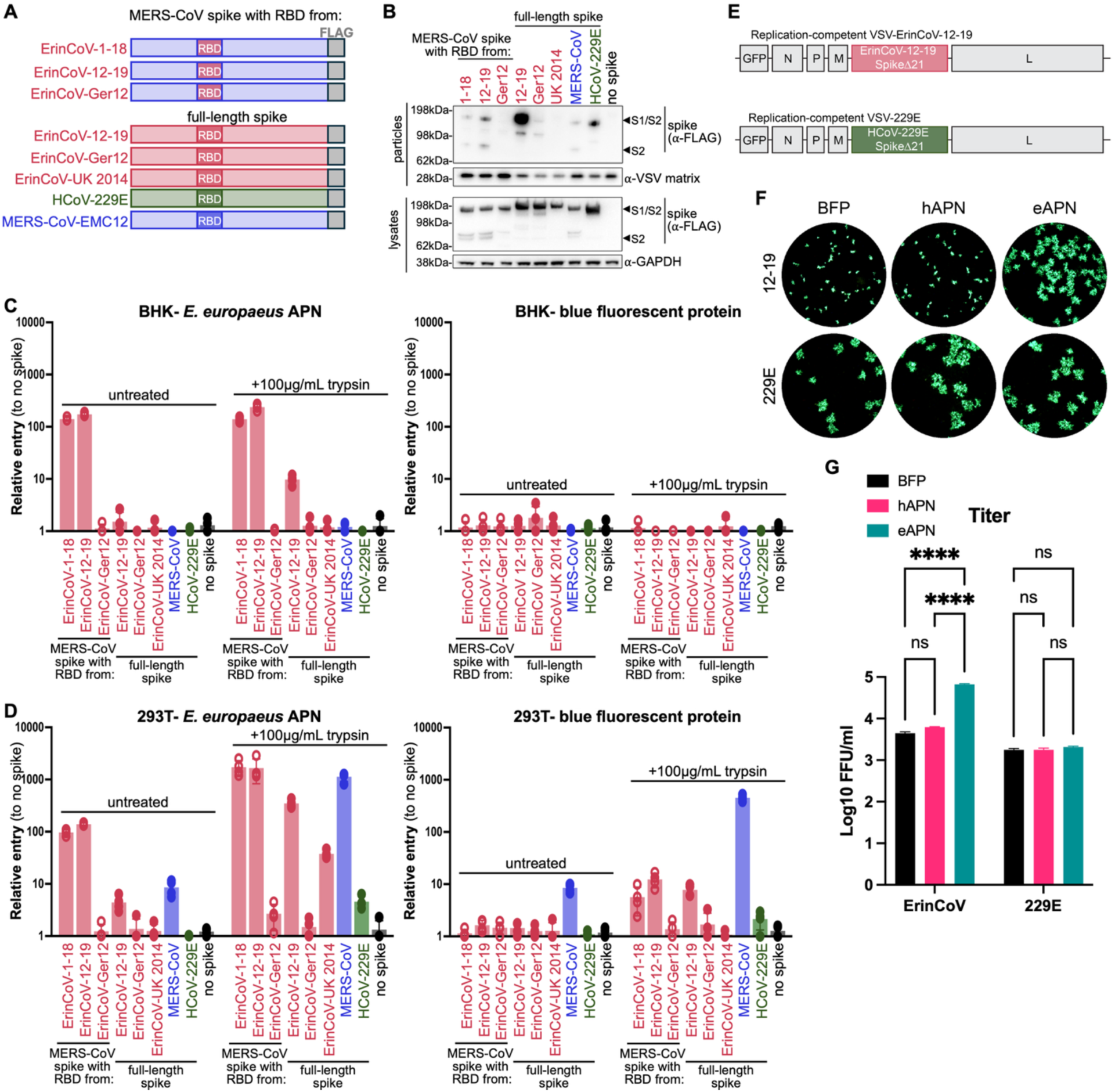
Full-length ErinCoV spike infects cells stably expressing hedgehog APN. **(A)** Schematic overview of wild-type and chimeric spikes used in this experiment **(B)** Western blot for spike and loading control proteins. **(C)** BHK cells or **(D)** 293T cells transduced to stably express either hedgehog APN or blue fluorescent protein (negative control) were infected with pseudotypes bearing either chimeric or wild-type spikes in quadruplicate. **(E)** Diagram of the replication-competent VSV (rcVSV) genome engineered to carry the spike glycoprotein of either ErinCoV-12-19 or HCoV-229E with C-terminal tail truncation. **(F)** Representative fluorescent focus-forming assay (FFA) images obtained after infection of Huh7.5 cells stably expressing the indicated APN ortholog with each rcVSV. **(G)** Quantification of infectious titers on the same Huh7.5-APN cell lines. Error bars show the mean +/- SEM of three independent experiments.

### Chimeric ErinCoV spikes reveal broader use of APN in ErinCoV RBDs

Due to the sequence similarity among hedgehog coronaviruses, we wondered if some of these viruses were truly incompatible with hedgehog APN as seen in our previous entry assay results (**Figure 1-2**). To further explore ErinCoV RBD compatibility with hedgehog APN, we generated chimeric ErinCoV spikes, whereby we exchanged the RBDs of the ErinCoV-12-19 spike and ErinCoV-Ger12 spikes with the other (**Figure 3A**). Spikes produced with an ErinCoV backbone are present in higher levels in concentrated pseudotypes than those produced in a MERS-CoV backbone, but this did not appear to correlate with viral entry (**Figure 3B, C**). Spikes produced with the ErinCoV-12-19 backbone appeared to have the highest rate of incorporation into viral particles and permitted viral entry with both ErinCoV RBDs. However, the higher incorporation of spikes produced with the ErinCoV-Ger12 backbone as opposed to those with the MERS-CoV backbone did not result in viral entry with either RBD. While the RBD from ErinCoV-12-19 permitted infection when incorporated in the MERS-CoV and ErinCoV-12-19 backbones, it did not when inserted into the ErinCoV-Ger12 backbone (**Figure 3C**). In contrast, the ErinCoV-Ger12 RBD only permitted viral entry when inserted into the ErinCoV-12-19 backbone. These results show that the ErinCoV-Ger12 RBD interacts with APN and can facilitate viral entry, despite being below the limit of detection in the initial assays.

**Figure 3.**
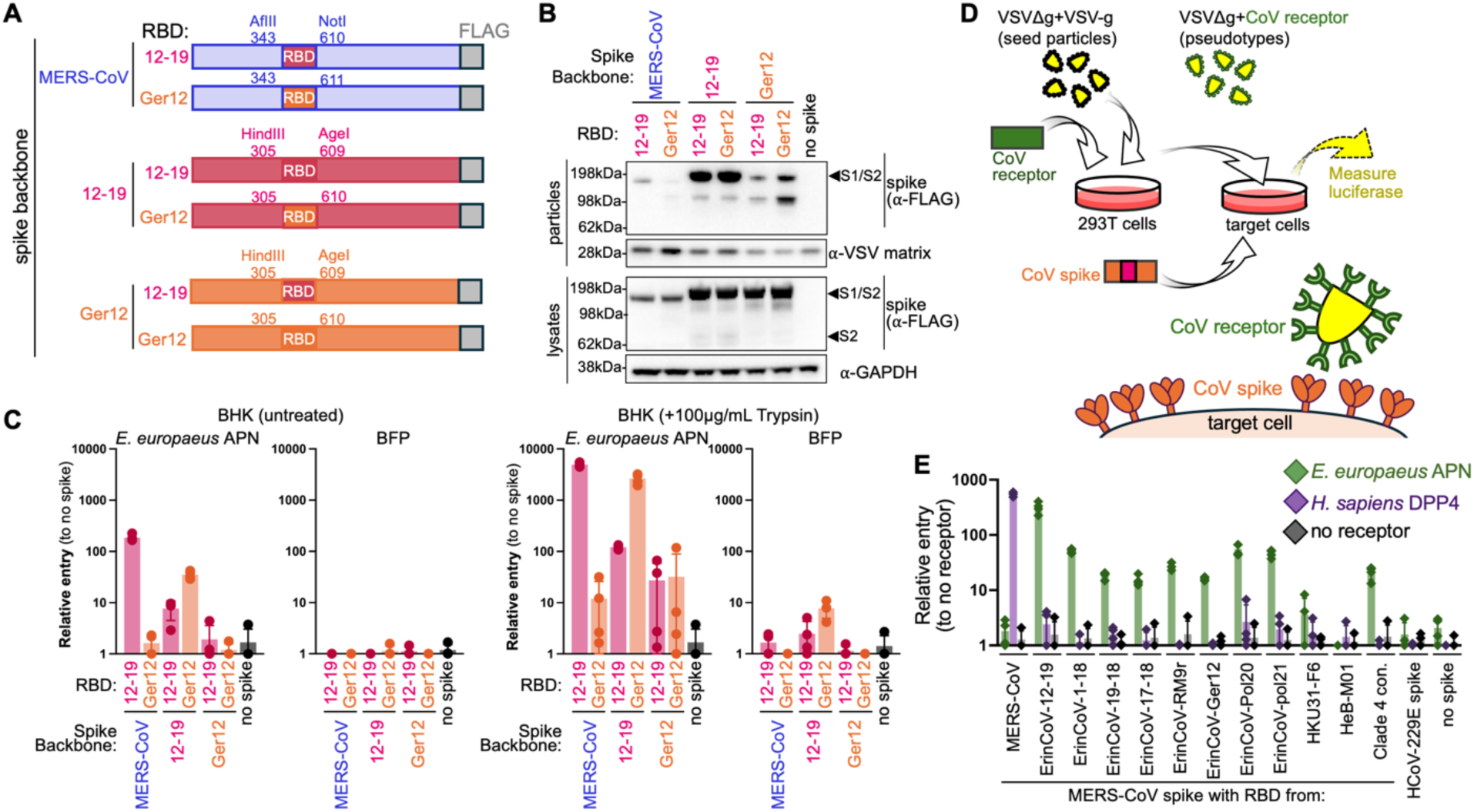
Alternative entry assays reveal cryptic ErinCoV receptor interactions. **(A)** Schematic overview of spikes used in this experiment **(B)** Western blot for FLAG (spike) and loading control protein expression **(C)** BHK cells transfected with hedgehog APN or BFP were infected with pseudotypes in the absence or presence of trypsin. **(D)** Overview of reverse entry assay with receptor based pseudotypes used to infect spike-expressing cells. **(E)** BHK cells transfected with indicated spikes were infected with pseudotypes bearing *E. europaeus* APN, human DPP4 or no receptor. Cells were infected in four technical replicates for all data presented; lines indicate standard deviation.

Given our results with the ErinCoV-Ger12 RBD (**Figure 3C**), we turned our attention to the other “non-viable” spikes from the initial experiments as well as an ErinCoV RBD consensus sequence (“Clade 4 Con.”)^52^. As an alternative way to assess the interaction between spike and receptor, we performed a reverse pseudotype assay and infected cells expressing viral spikes with VSV particles bearing viral receptors (**Figure 3D**). For this assay, we focused on the chimeric ErinCoV spikes from **Figure 1** that appeared non-viable with hedgehog APN. In line with our previous results with the ErinCoV-Ger12 RBD, the majority of the “non-viable” RBD chimeras from **Figure 1** mediated entry of hedgehog APN-pseudotyped VSV particles. Thus, our results demonstrate that all ErinCoVs found in European hedgehogs can interact with APN from European hedgehogs (**Figure 1, 3**).

### ErinCoV spikes interact with a limited number of species APN orthologues

In order to assess viral compatibility with other species we generated a panel of APN orthologues from 30 different species (**Figure 4**). BHK-21 cells were transfected with the APN orthologue panel with GFP serving as a negative control. Single-cycle VSV was pseudotyped using the wild-type ErinCoV-UK-2014, ErinCoV-Ger12, or ErinCoV-12-19 spikes, as well as chimeric MERS-CoV spikes with either the RBD from ErinCoV-Ger12 or ErinCoV-12-19. To enhance our ability to detect weak receptor interactions for this spike panel, we concentrated the viral pseudotypes 10-fold using Amicon Columns^71^. All the wild-type and chimeric spikes that we tested, including ErinCoV-Ger12, were capable of entering BHK cells expressing hedgehog APN (**Figure 4**), further confirming that viruses with an undetectable entry signal in our earlier assays also possess the capacity to use hedgehog APN as a receptor. The receptor specificity shown by the ErinCoV spikes was unexpectedly narrow, with only APN from the Cape elephant shrew (*Elephantulus edwardii*), brown rat (*Rattus norvegicus*), and domestic cat (*Felis catus)* supporting viral entry in addition to hedgehog APN (**Figure 4**).

**Figure 4.**
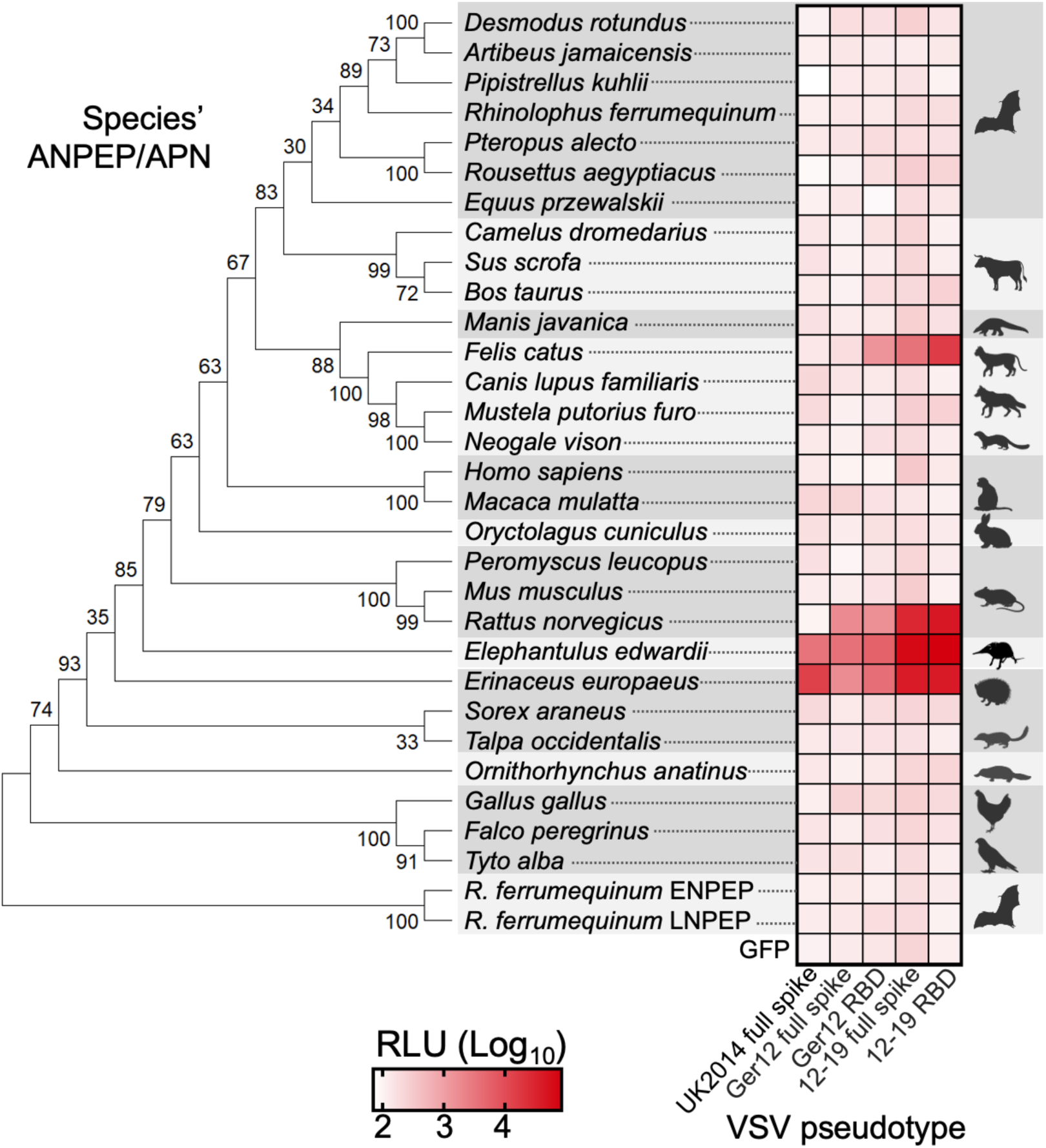
Cell entry screen with APN orthologues from different vertebrate species. Phylogenetic tree of amino acid sequences for alanyl glutamyl aminopeptidase (ANPEP), glutamyl aminopeptidase (ENPEP), and leucyl aminopeptidase (LNEP) from indicated species. Results from 1000 bootstraps are indicated over branches in the phylogenetic tree. BHK-21 cells transfected with the receptor orthologues and infected with pseudotypes bearing either wild-type ErinCoV spikes or chimeric MERS-CoV spikes with ErinCoV RBDs. Results shown are the mean of three technical replicates from two independent experiments.

### Structural basis for the interaction the between ErinCoV RBD and hedgehog APN

To characterize the molecular basis of the ErinCoV interaction with its host receptor, we determined the cryo-EM structures of the RBDs of both ErinCoV-12-19 and ErinCoV-Ger12 in complex with hedgehog APN. The complexes were determined in the presence of the APN inhibitor amastatin to promote the APN closed conformation (**Supplemental Figure 3**). The interactions between the bound amastatin and the protein are essentially identical to that observed in the human APN structure^72^, a reflection of the fact that the coordinating residues are identical in the two proteins. APN is a 2-fold symmetric dimer, and both complexes were determined with *C_2_* symmetry imposed, yielding global resolutions of 1.99 Å and 1.96 Å, respectively. Local refinement of the RBD and APN domains I, II, and III as a rigid body led to improved resolution and enhanced structural detail at the RBD-APN interface in both cases (local resolutions of 1.96 Å and 1.93 Å, respectively; **Supplemental Figure 4**, **Supplemental Table 1**). Due to the similarity in the sequences of the two ErinCoV RBDs, the complexes are nearly identical, differing by only a single amino acid in the RBD-APN interface as discussed below. In both complexes, an RBD is bound to each APN monomer through interactions involving only APN domain II (**Figure 5A**). Upon binding, the surface area buried on the ErinCoV RBD and the hedgehog APN range from 575 to 601 Å^2^, respectively, in the two complexes. Other alphacoronaviruses (HCoV-229E, CCoV-HuPn-2018, PRCV and TGEV) and porcine deltacoronavirus (PDCoV)^57–60^, have also been found to use APN as receptor and remarkably, the site on APN used by the ErinCoVs is distinct from that previously observed (**Supplemental Figure 5**).

**Figure 5.**
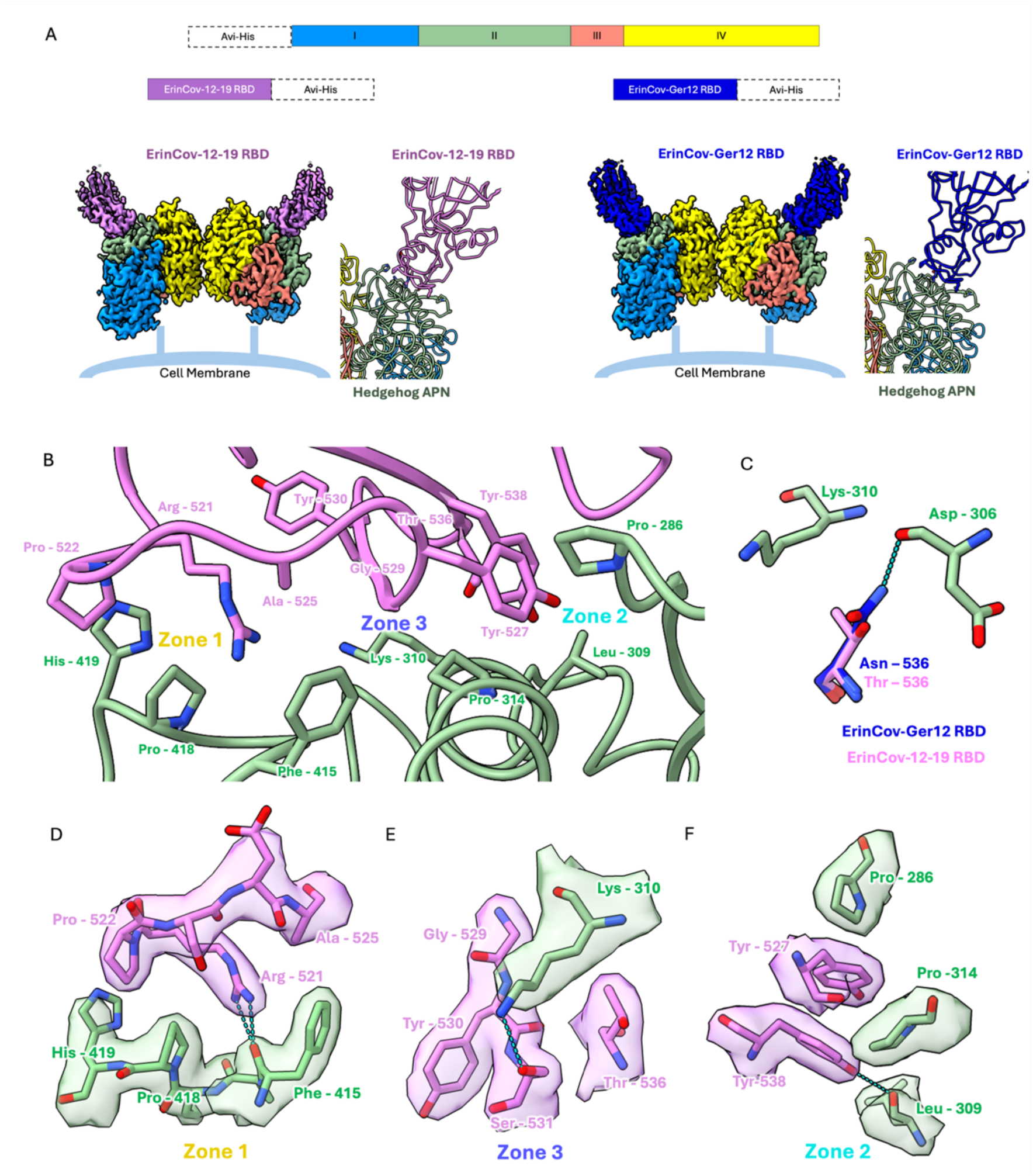
Hedgehog APN – ErinCoV RBD complex structures. **(A)** Domain representation of the hedgehog APN, ErinCoV-12-19 RBD and ErinCoV-Ger12 RBD expression constructs. The cryo-EM density maps of the ErinCoV-12-19 and ErinCoV-Ger12 RBDs in complex with hedgehog APN are colored by domain and depicted in their likely orientation relative to the cell membrane. The ribbon views show the RBD-APN (domain II) interface. In **(B)** and **(D-F)**, residues from the hedgehog APN and ErinCoV-12-19 RBD are colored violet and green, respectively. **(B)** Overview of the RBD-APN interface. **(C)** Stick models showing the only amino acid difference in the interfaces of the ErinCoV-Ger12 (blue) and ErinCoV-12-19 RBD (violet) complexes with hedgehog APN (green). In **(D-F)**, only density around the displayed residues (Zones 1, 2 and 3) from the locally refined cryo-EM map of the hedgehog APN – ErinCoV-12-19 RBD complex is shown. Hydrogen bonds are shown as dashed lines in cyan, in all cases.

The residues on the RBD (522-538) that mediate the interaction with APN are contained in the receptor binding subdomain on the same face that mediates the interaction between other merbecoviruses and their receptors (MERS-CoV-DPP4, HKU5-ACE2 and NeoCoV-ACE2; **Supplemental Figure 6**)^56,73,74^. The key residues mediating the ErinCoV-1219 RBD interaction with hedgehog APN can be grouped into three distinct zones (**Figure 5B**). Within Zone 1, apolar interactions are observed between RBD residue Pro-522 and APN residues Pro-418 and His-419. Apolar interactions are also observed between RBD residue Ala-525 and the side chain of APN residue Phe-415. The guanidino group of RBD residue Arg-521 also interacts with Phe-415, in this case through a cation-π interaction with its side chain and hydrogen bonds to its carbonyl oxygen (**Figure 5D**). The interactions in Zone 2 are centered on RBD residues Tyr-527 and Tyr-538. The former slots between APN residues Pro-286 and Pro-314 making apolar interactions with both, while the side chain of the latter is hydrogen bonded to the backbone carbonyl oxygen of APN residue Leu-309 (**Figure 5F**). These tyrosines interact with each other and are highly conserved among ErinCoV sequences, observations suggesting a key role in APN recognition. Zone 3 is defined by apolar interactions between RBD residues Thr-536, Gly-529 and Tyr-530 and the side chain of APN residue Lys-310 (**Figure 5E**). Additionally, the side chain nitrogen of Lys-310 is hydrogen bonded to the side chain hydroxyl of the RBD residue Ser-531. Thr-536 is replaced by Asn-536 in the ErinCoV-Ger12 RBD, and in its complex with APN, the amide nitrogen of the Asn-536 side chain is hydrogen bonded to the backbone carbonyl group of APN residue Asp-306 (**Figure 5C**). Notably, the chitobiose core of the *N-*glycan on APN residue Asn-362 stacks against the backbone of RBD residues Gln-533 and Gly-534 and this completes the interactions in Zone 3 (**Supplemental Figure 7**).

A biolayer interferometry-based assay was used to measure the affinity of the ErinCoV-12-19 and ErinCov-Ger12 RBDs to hedgehog APN and in both cases, measurements were made alternately with the RBD and APN coupled to the biosensor. Consistent with the high degree of structural similarity in the RBD-APN interface they both bound hedgehog APN with similar affinities (average *K_D_* values of ∼19 µM and ∼21 µM, respectively; **Figure 6**).

**Figure 6.**
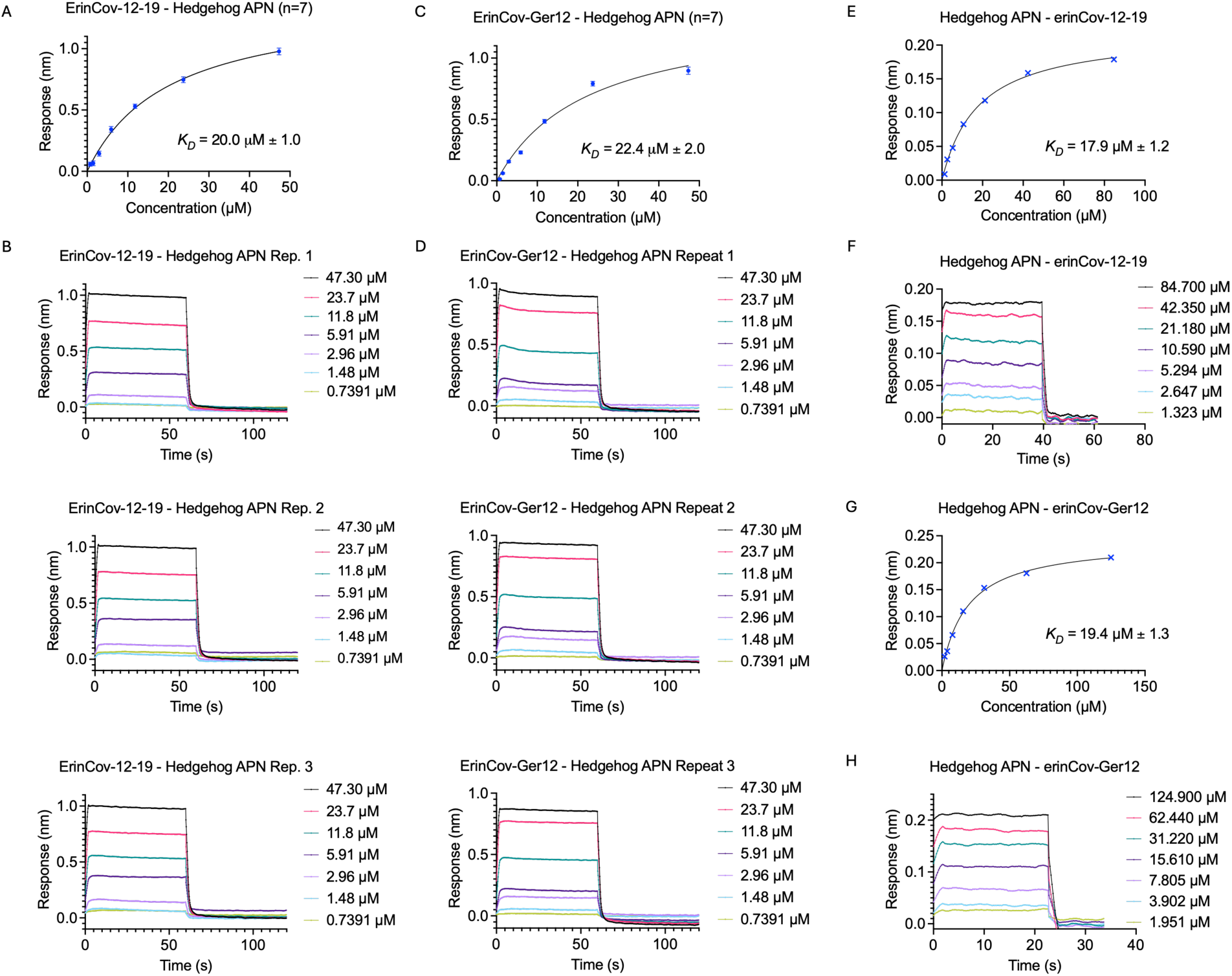
ErinCoV RBD binding affinity for hedgehog APN. **(A)** BLI steady state binding curves with the ErinCoV-12-19 RBD on the biosensor the hedgehog APN in solution. (n=7) **(B)** BLI binding sensorgrams with the ErinCoV-12-19 RBD on the biosensor and hedgehog APN in solution. Only three replicates are shown, and additional sensorgrams are shown in **Supplemental Figure 12**. **(C)** BLI steady state binding curves with the ErinCoV-Ger12 RBD on the biosensor and hedgehog APN in solution. (n=7) **(D)** BLI binding sensorgrams with the ErinCoV-Ger12 RBD on the biosensor and the hedgehog APN in solution. Only three replicates are shown, and additional sensorgrams are shown in **Supplemental Figure 12**. **(E)** BLI steady state binding curves with hedgehog APN on the biosensor and the ErinCoV-12-19 RBD in solution. **(F)** BLI binding sensorgrams with hedgehog APN on the biosensor and the ErinCoV-12-19 RBD in solution. **(G)** BLI steady state binding curves with hedgehog APN on the biosensor and the ErinCoV-Ger12 RBD in solution. **(H)** BLI binding sensorgrams with hedgehog APN on the biosensor and the ErinCoV-Ger12 RBD in solution.

### Mutations in the RBD and APN disrupt viral entry

To assess the importance of the residues found to interact in our hedgehog APN-RBD complexes, we repeated the entry assay after the introduction of mutations in either the RBD or APN. These include mutations in each of the three Zones described above (**Figure 7**). Point mutations in the ErinCoV-12-19 RBD within Zones 1 (Arg-521) and 2 (Tyr-527 and Tyr-538) reduced the entry signal for chimeric ErinCoV-12-19 to the level of the negative control in both BHK and 293T cells transfected with hedgehog APN (**Figure 7B**), and entry levels do not seem to be dependent on the level of spike incorporation in the pseudotype particles (**Figure 7C**). These results establish the importance of interactions in Zones 1 and 2 for ErinCoV entry.

**Figure 7.**
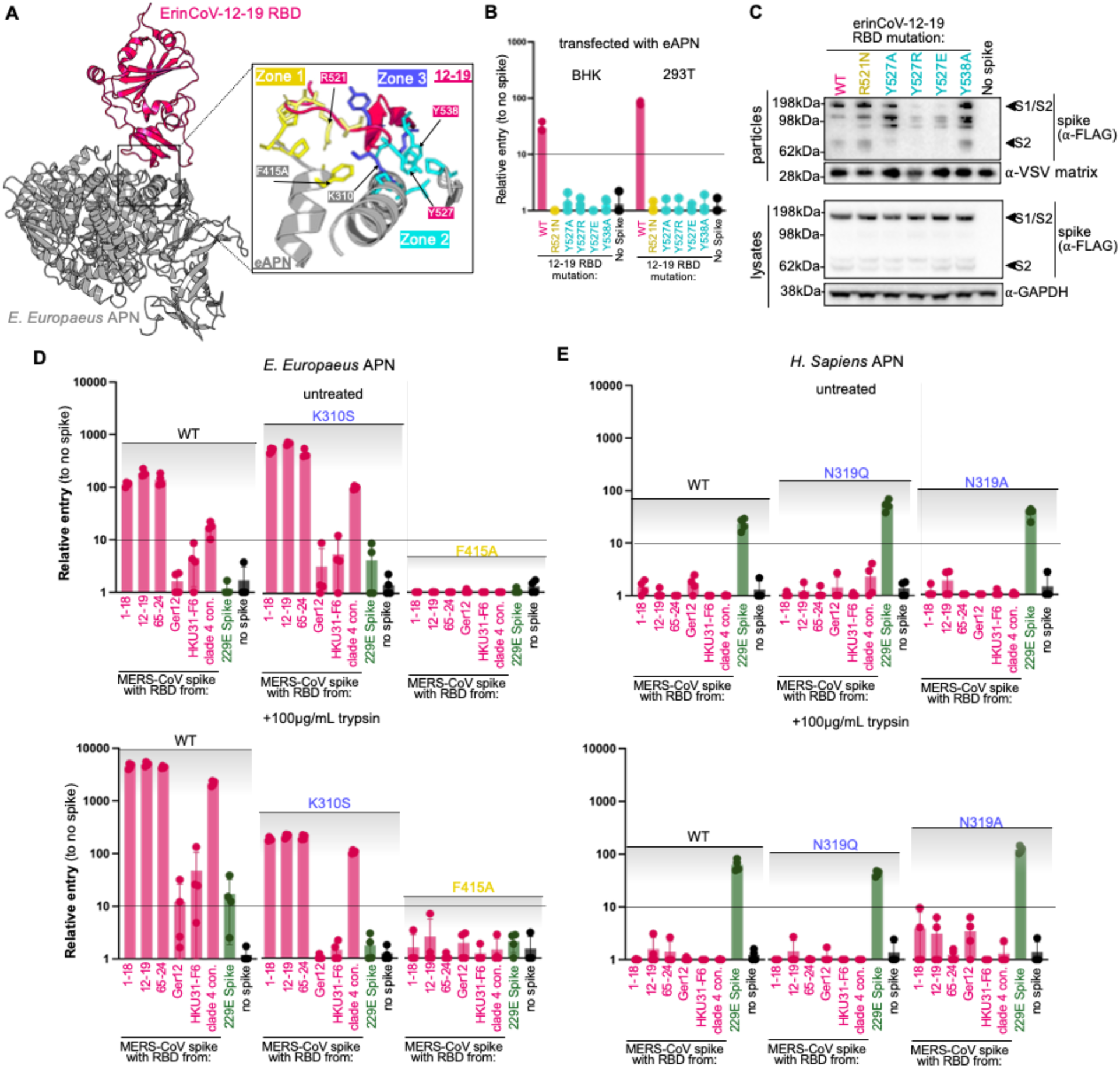
Structure-guided mutagenesis at the RBD:APN interface. **(A)** Overview of interaction between ErinCoV 12-19 RBD with hedgehog APN as determined by cryo-EM. **(B)** BHK or 293T cells were transfected with hedgehog APN and infected with pseudotypes bearing ErinCoV 12-19 RBD point mutations in the chimeric MERS-CoV spike backbone **(C)** Western blot for FLAG (spike) and loading control proteins in lysates and concentrated supernatants **(D)** BHK cells were transfected with hedgehog or **(E)** human APN mutants and infected with a small panel of ErinCoV RBD pseudotypes, in the presence or absence of trypsin. Cells were infected in four technical replicates

To test the effect of mutations in APN, viral entry was measured for pseudotypes containing RBDs that previously permitted viral entry: ErinCoV-1-18, −12-19, −65-24, and a consensus ErinCoV RBD, as well as ErinCoV-Ger12 and HKU31-F6. Mutating hedgehog APN residue Phe-425 to alanine reduced the entry signal for all the tested pseudotypes, further confirming the importance of Zone 1 interactions (**Figure 7D**). In contrast, mutating hedgehog APN residue Lys-310 in Zone 3 to a serine did not disrupt ErinCoV entry, an indication that the interactions with this residue are not critical for ErinCoV RBD recognition. Lys-310 is of particular interest as it corresponds to an *N*-glycosylated asparagine residue (Asn-319) in human APN and based on our structures, an *N*-glycan at this position would block the binding of ErinCoV to human APN. However, mutating Asn-319 to glutamine or alanine in human APN did not lead to ErinCoV entry (**Figure 7E**), an indication that differences at other locations are also important as discussed below. As expected, this mutation in human APN did not change its ability to mediate the entry of an HCoV-229E pseudotype (**Supplemental Figure 5**).

### APN sequence variation provides insight into host range

Our ErinCoV RBD-APN complexes have provided important insights into the APN structural features required for ErinCoV RBD binding and in conjunction with APN sequence data, a rationalization for the entry data shown in **Figure 4**. Perhaps most important is the absence of an *N*-glycan in the RBD-APN interface. We have identified APN residues Asp-306 and Lys-310 (hedgehog APN numbering) as positions where *N*-glycan sequons are found in APN sequences, and that would be expected to block ErinCoV binding if *N*-glycosylated (Type 1 difference, **Supplemental Figure 8A**). Only 8 of the 28 APN sequences tested lack *N*-glycan sequons at these positions and, as such, would have the potential to be bound by the ErinCoV RBD. Of these, four lack Pro-286 and/or Phe-415 (Type 2 difference **Supplemental Figure 8A**), APN residues found to make key interactions with the RBD in Zones 2 and 3. Of the remaining four APN sequences, the domestic dog APN (*Canis lupus familiaris*) has a glutamic acid residue at position Ala-283 (Type 3 difference, **Supplemental Figure 8A**) and this would create a steric conflict with the ErinCoV RBD residue Tyr-538, a residue shown to be critical for entry. This leaves only cat *(Felis catus),* brown rat (*Rattus norvegicus*) and elephant shrew (*Elephantuulus edwardii*) APNs from our panel which we have shown are the only APNs tested capable of mediating ErinCoV entry (**Figure 4**). This analysis provides a framework for assessing the likelihood that ErinCoV might bind one of the many APN sequences deposited from a wide range of species. Notably, human APN not only possesses an *N*-glycan at the equivalent of Lys-310 (Asn-319 in human) but in addition, Type 2 and Type 3 differences expected to negatively impact on binding (**Supplemental Figure 8B**).

## DISCUSSION

The cellular receptors used by many animal coronaviruses are unknown, hindering our ability to assess potential emergence of novel viral threats and prepare for future pandemics. The identification of merbecoviruses in high abundance in hedgehogs across Europe has raised concerns about their potential to infect humans. However, without knowing how ErinCoV infects host cells, scientists have been unable to study these viruses in controlled settings, confounding efforts to evaluate their zoonotic risk potential and develop countermeasures. Before the discovery of ACE2-dependent merbecoviruses like PDF2180 and NeoCoV, it was widely believed that merbecoviruses primarily used only DPP4 as a receptor. Now, multiple studies have shown that merbecovirus strains may use DPP4 or ACE2 as an entry receptor. Remarkably, a large phylogenetically related group of merbecoviruses found in hedgehogs use neither DPP4 nor ACE2 for cell entry. In this study, we demonstrate that merbecoviruses from European hedgehogs use aminopeptidase N (APN) from their host species—and a limited number of other species— as their receptor (**Figures 1 and 4**). Our structural analysis of the hedgehog RBD-APN complex demonstrates species barriers at the receptor binding level, which would likely prevent these viruses from spilling over into humans (**Figure 5, Supplemental Figure 8**).

APN was first identified as the receptor for Transmissible Gastroenteritis Coronavirus (TGEV), an alphacoronavirus that infects piglets^75^. It was also found to be the receptor for other alphacoronaviruses including the human coronavirus HCoV-229E^43^, porcine respiratory coronavirus (PRCV)^76^, the canine (CCoV) and feline (FIPV and FECV) coronaviruses^77^, and the canine-feline recombinant coronavirus (CCoV-HUPn-2018) that is also capable of infecting humans^59^. In addition to these alphacoronaviruses, the porcine deltacoronavirus (PDCoV) also uses APN as its receptor^57^. Our structural data show that the spike proteins of merbecoviruses identified in hedgehogs from geographically distinct regions utilize the same interface on hedgehog APN, but that this interface is distinct from those used by the others^57–60^ (**Figure 5; Supplemental Figure 5**). These observations suggest that during coronavirus evolution, APN has been independently selected multiple times to serve as the host receptor^58^. They also provide insight into the role of APN *N*-glycans in defining viral host range. As shown in **Supplemental Figure 5**, there are several APN *N*-glycans found on the distal face bound by these viruses and their locations vary among species. In the case of ErinCoVs (**Supplemental Figure 8**), any species with an *N*-glycan at a position equivalent to those within the ErinCoV-RBD–APN interface would likely be resistant to cross-species transmission. This convergence on different sites within the same receptor is analogous to what has been reported with coronavirus interactions with ACE2^78^. Taken together, the remarkably limited number of identified coronavirus receptors reflects the convergent evolution of their entry mechanisms.

Interestingly, while our data suggests that most if not all ErinCoVs use APN as a receptor, the spikes and RBDs from some viruses performed poorly and did not produce as clear an entry signal as others, except when the spike backbone was changed or the viral pseudotype assay was reversed (**Figure 3**). For example, ErinCoV-12-19 clearly infected cells in all our assays if the wild-type spike or the chimeric spike containing the 12-19 RBD in the MERS-CoV backbone was used (**Figure1-3)**. On the other hand, ErinCoV-Ger12 did not enter cells in most of the assays, except when the pseudotypes were concentrated 10-fold (**Figure 4**), the backbone was changed from the MERS-CoV to the ErinCoV-12-19 spike, or if the entry assay was reversed (**Figure 3**). In support of these results, our binding data shows that the affinity of the ErinCoV-Ger12 RBD is similar to or slightly weaker than that of the ErinCoV-12-19 RBD for hedgehog APN (**Figure 6**). Moreover, our structural analysis shows that the ErinCoV-Ger12 and ErinCoV-12-19 RBD complexes with hedgehog APN are essentially identical (**Figure 5**). Notably, both MERS-CoV spike-based chimeric and wild-type ErinCoV-Ger-12 spikes both failed to enter cells (**Figure 2**). Therefore, the failure to detect entry for both the wild-type and MERS-CoV spike-based chimeric spikes may be the result of a yet unknown species barrier. The reverse entry assay we employed to detect entry of “non-viable” chimeric spikes from **Figure 1** may have allowed for turnover of cell-produced membrane proteins, including the viral spike gene (**Figure 4D, E**). This membrane turnover may have allowed for destabilized spikes to be replaced with freshly-produced trimers, which would be advantageous over spike-based pseudotypes, which have no metabolism and cannot undergo membrane turnover. It is possible that the infection conditions used in our assays are not suitable for some ErinCoVs, which may be more adapted toward hedgehog cell physiology. Our results from this study suggest multiple barriers for viral entry exist for the ErinCoVs, including temperature (**Supplemental Figure 2**), APN sequence variation across species (**Figure 3; Supplemental Figure 8**), and possibly host protease compatibility (**Figure 1-3**). Future studies with hedgehog derived cell cultures may help elucidate additional species barriers.

While inconclusive, our entry results with ErinCoVs found in Chinese hedgehogs (HKU31-HeB-M01, HKU31-F6) were not completely negative (**Figure 1E-G**, **Figure 3E**). In light of our other results showing discrepancies between ErinCoV-Ger12 entry and binding, it may be possible that these other ErinCoVs also bind hedgehog APN. Our earlier study identifying the receptor for HKU5 showed high-species specificity between virus and host, suggesting that ErinCoVs found in *Erinaceus amurensis* may use APN from this species but few others^52^. Unfortunately, the genome for *Erinaceus amurensis* is not available, preventing us from testing the APN orthologue from this species.

Our data suggests ErinCoVs are likely to have limited species tropism due to several potential barriers at the level of entry. With these results in mind, the hedgehog strains studied herein face multiple obstacles for efficient cross species transmission and/or replication and likely pose reduced risk towards humans. However, the enormous quasispecies diversity in the merbecovirus subgenus, coupled with the potential to recombine and exchange genetic elements, serves as a warning that newly identified viruses can emerge suddenly and change this understanding. For example, while HKU5 viruses exhibit limited usage for ACE2 of other species, a second lineage of HKU5 viruses was recently reported with broader ACE2 compatibility, and the recent spill-over of HKU5 from bats to mink resulted in a virus with far enhanced human ACE2-binding potential^13,78,79^. We show here that, similar to HKU5, ErinCoVs possess compatibility with limited species but are compatible with cat, rat, and elephant shrew APN. Taken together, the possibility remains that ErinCoVs may spillover to a species that is in proximity with humans, further adapt, and spillover to humans. In general, broader antiviral interventions are urgently needed to prevent future pandemics.

## METHODS

### Cells

No commonly misidentified cells were employed for this study. 293T (ATCC CRL-3216) and BHK-21 (ATCC CCL-10) cells were propagated in DMEM (Gibco) supplemented with 10% FBS (GE HyClone), penicillin-streptomycin (Gibco), and L-glutamine (Gibco) under 5% CO_2_ at 37°C. Cell line species origins were confirmed by cytochrome sequencing and all cell lines were verified as mycoplasma negative with MycoSniff PCR Kit (MP Bio).

### Expression and purification of hedgehog APN and the ErinCoV spike RBDs

Soluble secreted forms of hedgehog APN (NCBI accession XP_007516437.1) and the RBDs of the ErinCoV-12-19 (NCBI accession: MW246800) and ErinCoV-Ger12 (NCBI accession KC545383) spike proteins were produced in HEK293F cells and purified as previously described^69^. The cDNA of the hedgehog APN (52-960) was codon-optimized and N-terminal Avi and 6-His tags were incorporated. For the ErinCoV-12-19 RBD (364-572) and the ErinCoV-Ger12 RBD (364-573), the cDNA was codon-optimized and a linker containing an *N*-glycosylation site (SGAPNSTSAP) was added to the C-terminus followed by 6-His and Avi tags. In all cases, the secreted proteins were purified by Ni^2+^-NTA affinity chromatography followed by gel filtration (Superdex 200 Increase, Cytiva). Amino acid sequences for all the constructs can be found in **Supplemental Figure 9**.

### Biolayer Interferometry (BLI) binding studies

The hedgehog APN protein was biotinylated using NHS-PEG4-Biotin (ThermoFisher, Catalogue Number: A39259) according to the manufacturer’s instructions. NHS-PEG4-biotin was dissolved in DMSO to produce a stock solution at a concentration of 1 mM. 2 µl of the stock solution was added to 300 µl of the hedgehog APN solution (1 mg/ml) in PBS, pH 7.4. The reaction mixture was incubated at room temperature for 120 mins, followed by removal of excess NHS-PEG4-biotin by gel filtration chromatography (Superose 6 Increase, Cytiva) in PBS, pH 7.4. The biotinylated hedgehog APN was immobilized onto streptavidin biosensors (Octet Streptavidin Biosensor, Sartorius) to give up to a 2 nm response. The D1 proteins were prepared and serially diluted in kinetics buffer (PBS, pH 7.4, 0.1% Tween 20) for the binding study. A reciprocal experiment with the ErinCoV RBDs coupled to the biosensor was performed in the same way. Response versus concentration was plotted at the steady state (10 s after reaching the steady state plateau), and the *K_D_* was determined by fitting to a 1:1 model using GraphPad Prism.

### Cryo-EM Data Collection

Holey gold film-coated EM grids with 2 µm holes were prepared as previously described^80^. A 1 mM stock solution of the APN inhibitor amastatin (Thermo Fisher Catalogue Number: A1276) was prepared in water. The hedgehog APN and both ErinCoV RBD solutions were concentrated and re-purified by gel filtration (Superdex 200 Increase, Cytiva) in 10 mM Tris-HCl buffer, pH 7.5 containing 50 mM NaCl just prior to grid vitrification. For both the ErinCoV-12-19 and ErinCoV-Ger12 RBD complexes with hedgehog APN, the sample solution was prepared by mixing 10 µL of a 1.5 mg/mL APN solution, 5 µL of a 1 mg/mL RBD solution and 0.5 µL of the 1 mM amastatin solution. The resulting samples were incubated for 30 min at room temperature before grid vitrification using a Vitrobot Mark IV device.

Grid screening and optimization was performed on a Thermo Fisher Scientific Glacios 2, 200 kV microscope equipped with a Falcon 4i direct electron detector. High-resolution data collection was performed on a Thermo Fisher Scientific Titan Krios G3, 300 kV microscope equipped with a Falcon 4i direct electron detector. For both data sets, movies were collected at 165,000 × nominal magnification, resulting in a calibrated pixel size of 0.73 Å. Each movie was recorded in counting mode with a 4 s exposure time and saved in EER format. The exposure rate was adjusted to 7 electrons per pixel per second, resulting in a total exposure of 53 electrons per Å^2^. The data were collected with a 1,000 nm to 2,000 nm defocus. A total of 9,223 movies were collected for the hedgehog APN – ErinCoV-12-19 RBD dataset, and 9,493 movies were collected for the hedgehog APN – ErinCoV-Ger12 RBD dataset.

### Cryo-EM Data Processing **(Supplemental Figure 10, Supplemental Figure 11)**

Both datasets were processed using the following procedures. All data analysis steps were performed with cryoSPARC 4.6.0^81^. Both datasets were analyzed without super-resolution mode. Movies were aligned with patch-based motion correction, and contrast transfer function (CTF) parameters were estimated with patch-based CTF estimation. Particle selection was done with Topaz. The Topaz model for particle selection was generated from 2D classification jobs of particle images selected by automatic picking with a Gaussian blob from 200 randomly selected micrographs. Multiple rounds of 2D and 3D classification were used to remove low-quality particles from the Topaz-picked particles. For the hedgehog APN – ErinCoV-12-19 RBD complex, the final dataset included 1,209,700 particles. For the hedgehog APN – ErinCoV-Ger12 RBD complex, the final dataset included 971,683 particles. Further improvement in CTF parameter estimation and motion correction was achieved by performing global CTF refinement and local motion correction. Homogenous refinement was carried out with *C_2_* symmetry imposed, yielding a global resolution of 1.99 Å for the hedgehog APN – ErinCoV-12-19 RBD complex and 1.96 Å for the hedgehog APN – ErinCoV-Ger12 RBD complex. To improve the resolution in the vicinity of the RBD–APN interface, further 3D classification was carried out for symmetry expanded particles (with *C_2_* symmetry) using a mask containing APN domain I, domain II and domain III and the RBD. Particles with no bound RBD were removed from the particle stack, resulting in a final particle stack (with symmetry expansion) of 2,244,653 particles for the hedgehog APN-ErinCoV-1219 RBD complex and 1,693,136 particles for the hedgehog APN – ErinCoV-Ger12 RBD complex. Local refinement with the same mask used for 3D classification was carried out, yielding a map with a global resolution of 1.96 Å for the hedgehog APN – ErinCoV-12-19 RBD complex and 1.93 Å for the hedgehog APN – ErinCoV-Ger12 RBD complex.

### Model Building and Refinement

The hedgehog APN – ErinCov-12-19 RBD complex model (APN monomer and associated RBD) was built manually into the map using UCSF ChimeraX^82^ and Coot^83^. Initial models for the APN monomer and the RBD were generated using the AlphaFold online server^84^. The polypeptide chains could be traced from residues 61-960 for the APN monomer and residues 365-572 for the RBD. The model was refined by an iterative process involving model building in Coot, molecular dynamics-assisted refinement using the ISOLDE tool^85^ in UCSF ChimeraX and Real-Space refinement in Phenix^86^. 12 *N*-glycans were also built and refined using Coot and ISOLDE. The models for the hedgehog APN – ErinCov-12-19 RBD dimer, the hedgehog APN – ErinCoV-Ger12 RBD monomer and the hedgehog APN – ErinCoV-Ger12 RBD dimer were built and refined in a similar way using the hedgehog APN – ErinCoV-12-19 RBD model as a reference. All the models were validated using the comprehensive validation (cryo-EM) tool in Phenix. The molecular graphics were generated in UCSF ChimeraX^87^.

### Erinaceus europaeus APN cell lines

APN from *Erinaceus europaeus* was sub-cloned into a lentiviral expression vector with PCR and used to produce lentiviral particles in 293T cells^88,89^. Supernatants were filtered, aliquoted and frozen at −80°C. BHK cells and 293T cells were transduced with lentiviral expression vectors expressing either receptor orthologues or Blue Fluorescent Protein as a negative control. Cells were maintained for 3 passages under selective pressure before use in experiments and were maintained under selection with 1μg/mL puromycin (Fisher; BHK and 293T) during the entire study.

### Phylogenetic Analyses

For phylogenetic analyses of APN sequences, amino acid sequences were aligned using MAFFT algorithm (version 7.526). Phylogenetic trees were then constructed using the Maximum Likelihood method with 1000 bootstrap replicates in MEGA11 (version 11.0.73). The optimal protein model and parameters for the phylogenetic analysis were selected based on recommendations provided by the MEGA11 software. For phylogenetic analysis of merbecovirus RBDs, amino acids were aligned using Clustal multiple sequence alignment software. A maximum likelihood phylogenetic tree was produced with PhyML v. 3.0^52,90,91^, and cladograms were produced using FigTree v1.4.4 (https://github.com/rambaut/figtree).

### Plasmids

As described previously, MERS-CoV/EMC12 spike (accession number JX869059) was codon optimized, C-terminally FLAG tagged, and silent mutations were introduced near spike amino acids 341 and 617 to generate AflII and NotI restriction digest sites. RBDs were codon optimized for human cells, synthesized (IDT DNA), and used in downstream Infusion based cloning (Takara Bio) reactions to assemble full-length chimeric spike proteins. A complete list of accession numbers for the spike panel can be found in **Supplemental Figure 1**. *Erinaceus europaeus* ACE2, transcript variant 1 (GenBank XM_060183012.1), DPP4 (GenBank XM_060177812.1) and APN (GenBank XM_007516375.3) were synthesized (IDT DNA) and cloned into pcDNA3.1+ with NheI and ApaI sites. Sequence data for the 29 alanyl glutamyl aminopeptidase (ANPEP), 1 glutamyl aminopeptidase (ENPEP), and 1 leucyl aminopeptidase (LNPEP) receptors were obtained from GenBank and are detailed in **Table 1**. All receptor and viral gene sequences utilized in this study were synthesized by either GenScript or IDT DNA. Human codon-optimized sequences of ANPEP, ENPEP, and LNPEP, each tagged with a C-terminal HA-tag, were synthesized and subcloned into the pUC57 expression vector using the BsmBI restriction site. Additionally, human codon-optimized sequences of hedgehog coronavirus (CoV) spike proteins, along with chimeric spikes generated by replacing the MERS-CoV RBD with either the ErinCoV-Ger12 RBD or the ErinCoV-12-19 RBD—both tagged with a C-terminal Flag tag, were synthesized and subcloned into expression vectors.

**Table 1.** List of the ANPEP, ENPEP, and LNPEP orthologs used in this study.

| Gene | Species | NCBI Reference Sequence |
| --- | --- | --- |
| ANPEP | *Homo sapiens* | NP_001141.2 |
| ANPEP | *Mus musculus* | NP_032512.2 |
| ANPEP | *Rattus norvegicus* | NP_112274.1 |
| ANPEP | *Gallus gallus* | NP_990192.2 |
| ANPEP | *Sus scrofa* | XP_020953963.1 |
| ANPEP | *Canis lupus familiaris* | NP_001139506.1 |
| ANPEP | *Bos taurus* | NP_001068612.1 |
| ANPEP | *Felis catus* | NP_001009252.2 |
| ANPEP | *Macaca mulatta* | XP_001093727.3 |
| ANPEP | *Oryctolagus cuniculus* | NP_001075795.1 |
| ANPEP | *Ornithorhynchus anatinus* | XP_028907451.1 |
| ANPEP | *Sorex araneus* | XP_004617569.1 |
| ANPEP | *Mustela putorius furo* | XP_012917463.1 |
| ANPEP | *Falco peregrinus* | XP_005239470.3 |
| ANPEP | *Elephantulus edwardii* | XP_006885282.1 |
| ANPEP | *Pteropus alecto* | XP_006905412.1 |
| ANPEP | *Erinaceus europaeus* | XP_007516437.1 |
| ANPEP | *Equus przewalskii* | XP_008508865.1 |
| ANPEP | *Camelus dromedarius* | XP_031296423.1 |
| ANPEP | *Rousettus aegyptiacus* | XP_016007055.1 |
| ANPEP | *Manis javanica* | XP_017506854.2 |
| ANPEP | *Desmodus rotundus* | XP_024417425.1 |
| ANPEP | *Peromyscus leucopus* | XP_028729561.1 |
| ANPEP | *Tyto alba* | XP_042660314.1 |
| ANPEP | *Rhinolophus ferrumequinum* | XP_032956109.1 |
| ANPEP | *Pipistrellus kuhlii* | XP_036271433.1 |
| ANPEP | *Artibeus jamaicensis* | XP_037000911.2 |
| ANPEP | *Talpa occidentalis* | XP_037365785.1 |
| ANPEP | *Neogale vison* | XP_044086346.1 |
| ENPEP | *Rhinolophus ferrumequinum* | XP_032962348.1 |
| LNPEP | *Rhinolophus ferrumequinum* | XP_032967111.1 |

### Pseudotype production

For ErinCoV panels, pseudotypes were generated in a manner based on previous protocols^52,64,67,92^. 293T cells were seeded in poly-lysine treated 6-well plates and incubated at 37°C for 24 hours. Cells were then transfected with spike plasmids using polyethyleneimine (Poly Sciences) and moved to a 34°C incubator for the remainder of preparation. 24-hours post transfection, cells were infected with VSV-ΔG-pseudotype VSV particles at a multiplicity of infection of two in serum-free media and left to incubate for one hour. To remove seed particles, cells were then washed three times and left in serum free media. 24-hours post seed particle infection, media was removed from the cells and replaced with fresh serum-free media. Pseudotypes were collected at 48-hours post infection, clarified by centrifugation, aliquoted, and stored at −80°C until needed.

For the APN panel, pseudotyped VSV particles were generated as following established protocols^93^. Briefly, HEK-293T cells were transfected with expression vectors encoding various coronavirus spike proteins. After 24 hours, these cells were infected with a replication-deficient VSV vector lacking the glycoprotein gene (VSV-ΔG) and containing Renilla luciferase reporter gene. After a 1-hour incubation at 37°C, the inoculum was removed, and the cells were washed twice with 1X PBS. To neutralize the residual input virus, media containing the anti-VSV-G antibody clone I4 (Kerafast) was then added. Supernatant containing pseudotyped viruses were collected at 24- and 48-hours post-infection, clarified by centrifugation, and passed through a 0.45 μm PVDF syringe filter (MilliporeSigma) to remove cellular debris. The harvested VSV-ΔG-pseudoviruses were then concentrated 10-fold using 100 kDa Amicon™ Ultra-15 Centrifugal Filter Units (MilliporeSigma) and stored at −80°C until further use.

### Pseudotype entry assays

For ErinCoV panels, entry assays were performed using standard approaches^52,65,66^. For experiments where receptors were transfected, 293T or BHK cells were seeded in black 96-well plates, transfected with receptor plasmids the next day, and infected 24 hours later with equal volumes of viral pseudotypes. For validation of the initial large ErinCoV pseudotype panel and full spike experiments, 293T or BHK cells stably transduced with *Erinaceus europaeus* APN were seeded in black 96-well plates and subsequently infected 24 hours later. Infections were conducted on ice to prevent unintended, premature trypsin activity. Pseudotypes were diluted with either HBSS or HBSS with trypsin (not TPCK-treated) at a final concentration of 100μg/mL trypsin. Cells were washed once with cold PBS prior to infection with pseudotypes, centrifuged at 4°C, 1200 x g for 1 hour and incubated at 37°C overnight. Luciferase was measured using Bright-Glo reagent (Promega) approximately 18 hours post-infection. Relative entry was calculated by dividing raw luciferase values for each spike by the signal for no-spike pseudotypes. Plates were each measured 4 times to reduce error generated by plate reader background, analyzed individually and then averaged across all four measurements. Relative entry values for all four replicates were averaged and plotted as heatmaps using Microsoft Excel.

For the APN panel, BHK-21 cells were transfected with plasmids encoding various animal receptors (ANPEP, ENPEP, and LNPEP) using PEI MAX (Polysciences). The following day, 50,000 receptor-expressing cells were seeded into black-walled, clear-bottom 96-well plates in 100 µl of medium. VSV pseudoviruses bearing different coronavirus spike proteins and carrying the Renilla luciferase (RLuc) reporter gene were then added at a 1:1 volume ratio in DMEM supplemented with 2% fetal bovine serum (FBS), and the plates were centrifuged at 1200 g for 1 hour at 4 °C. After 24-hour incubation, the cells were lysed to measure luciferase activity. To do this, 25 μl of 1X Renilla lysis buffer (Promega), prepared by diluting 5X buffer with phosphate-buffered saline (PBS), was added to each well and incubated for 20 minutes at room temperature with shaking. Then, 25 μl of Renilla luciferase substrate, diluted at 1:100 in assay buffer, was added to the lysates. Luminescence was measured in triplicate using a BioTek Synergy microplate reader with Gen5 software, and the relative light units (RLU) were normalized and analyzed using GraphPad Prism (v10.2).

### Replication-competent VSV

Replication-competent VSV (rcVSV) ErinCoV-12-19 and VSV-229E with deletion of the 21 C-terminal amino acids expressing green fluorescent protein (GFP) were generated as previously described^94^. Briefly, HEK293T cells stably expressing T7 RNA polymerase were transfected with plasmids encoding VSV N (Addgene #64087), P (Addgene #64088), L (Addgene #64085), G (Addgene #64084), and an antigenome copy of the viral genome (Addgene #31842) under control of the T7 promoter. Rescue supernatants were collected 24 hours-post transfection and clarified by centrifugation (5 minutes at 500 x g). Virus clones were plaque-purified and subsequently amplified on Huh-7.5 cells expressing cognate APN receptors at 37°C with 5% CO2. Viral supernatants were harvested after extensive cytopathic effects were observed and clarified by centrifugation (5 minutes at 500 x g). Aliquots were maintained at −80°C.

### Fluorescent focus forming assay

Huh-7.5 cells stably expressing the cognate APN receptor were seeded at 2.5×10^4^ cells per well in black, clear-bottom 96-well plates 24 hours before infection. Replication-competent VSV (rcVSV) stocks were serially diluted 10-fold in PBS in round-bottom 96-well plates, then transferred to the cell plates in duplicate and allowed to adsorb for 1 hour at 37°C, 5% CO_2_. After adsorption, an overlay of Opti-MEM supplemented with 1% carboxymethylcellulose (CMC) was added, and plates were incubated for 72 hours. Cells were then fixed with 10% neutral-buffered formalin for 1 hour at room temperature, rinsed with distilled water, and air-dried. Fluorescent foci were imaged and quantified using a BioSpot reader.

### Statistics and reproducibility

Cell entry data shown in **Figures 1**, **2**, **4**, and **7** is from four technical replicates, but representative of multiple biological replicates performed over the course of this study. Species APN screening results in **Figure 3** are from 3 technical replicates and 2 biological replicates. Between 3 and 4 technical replicates were chosen to maximize statistical relevance, while still allowing for scalable experiments. Throughout this study, experiments were performed by different labs, with different batches of pseudotypes, different batches of cells and at different times.

### Western blots

As described previously^52,64^, supernatants were removed from 293T cells transfected with spike plasmids, and the cells were lysed in 1% SDS lysis buffer and stored at −80°C until use. Pseudotyped particles were concentrated from producer cell supernatants that were overlaid on a 15% Opti-Prep cushion in PBS (Sigma) and centrifuged at 20,000× g for 2 hours at 20°C. Lysates and concentrated particles were clarified, reduced, and boiled. Proteins were separated using 10% Bis-Tris PAGE gel (Thermo Fisher). After protein transfer, membranes were probed for FLAG, GAPDH and/or VSV-m expression and imaged using SipersSignal West Substrate (Thermo Fisher) and a digitial developer (BioRad)^52^.

### Data availability

Accession numbers for all spike sequences used in this study can be found in **supplemental Figure 1** and accession numbers for all APN sequences used are listed in **Table 1**. Data generated from cryo-EM studies of the ErinCoV-12-19 and ErinCoV-Ger12 RBDs have been deposited at the Protein Data Bank (PDB) and Electron Microscopy Data Bank (EMDB) under accession codes PDB: 9OW3, 9OW4, 9OW5 and 9OW6 and EMD-70927, EMD-70928, EMD-70929, and EMD-70930, respectively.

## SUPPLEMENTAL INFORMATION

**Supplemental Table 1:** Cryo-EM data collection/refinement and PDB model statistics.

| Structure | Hedgehog APN -<br>ErinCov-12-19 | Hedgehog APN -<br>ErinCov-12-19<br>(Local Refine) | Hedgehog APN -<br>ErinCov-Ger12 | Hedgehog APN -<br>ErinCov-Ger12<br>(Local refine) |
| --- | --- | --- | --- | --- |
| **Data Collection** |  |  |  |  |
| Electron Microscope | Titan Krios G3 | Titan Krios G3 | Titan Krios G3 | Titan Krios G3 |
| Camera | Falcon 4i | Falcon 4i | Falcon 4i | Falcon 4i |
| Voltage (kV) | 300 | 300 | 300 | 300 |
| Nominal Magnification | 165,000 | 165,000 | 165,000 | 165,000 |
| Calibrated physical pixel size (Å) | 0.73 | 0.73 | 0.73 | 0.73 |
| Total exposure (e/Å<sup>2</sup>) | 53 | 53 | 53 | 53 |
| Exposure rate (e/pixel/s) | 7 | 7 | 7 | 7 |
| Exposure time (s) | 4 | 4 | 4 | 4 |
| File format | EER | EER | EER | EER |
| Defocus range (µm) | 1.0 to 2.0 | 1.0 to 2.0 | 1.0 to 2.0 | 1.0 to 2.0 |
| **Image Processing** |  |  |  |  |
| Super-resolution mode | No | No | No | No |
| Particle images contributing to maps | 1,209,700 | 2,244,653 | 971,683 | 1,693,136 |
| Applied symmetry | *C*<sub>2</sub> | *C*<sub>1</sub> | *C*<sub>2</sub> | *C*<sub>1</sub> |
| Map global resolution (FSC = 0.143, Å) | 1.99 | 1.96 | 1.96 | 1.93 |
| **Model building** |  |  |  |  |
| Modeling software | COOT,PHENIX,<br>ISOLDE | COOT,PHENIX,<br>ISOLDE | COOT,PHENIX,<br>ISOLDE | COOT,PHENIX,<br>ISOLDE |
| RMSD bond length (Å) | 0.013 | 0.013 | 0.013 | 0.013 |
| RMSD bond angle (°) | 1.675 | 1.679 | 1.684 | 1.697 |
| Ramachandran outliers (%) | 0.00 | 0.00 | 0.00 | 0.00 |
| Ramachandran favoured (%) | 96.65 | 96.92 | 97.11 | 96.93 |
| Rotamer Outlier (%) | 0.31 | 0.10 | 0.20 | 0.20 |
| Clashscore | 0.39 | 0.00 | 0.06 | 0.06 |
| MolProbity score | 0.85 | 0.68 | 0.68 | 0.71 |
| **Map to model fit** |  |  |  |  |
| CC (mask) | 0.88 | 0.89 | 0.89 | 0.90 |
| CC (box) | 0.84 | 0.80 | 0.84 | 0.78 |
| CC (peaks) | 0.77 | 0.72 | 0.77 | 0.70 |
| CC (volume) | 0.88 | 0.89 | 0.89 | 0.90 |
| Protein residue number | 2226 | 1112 | 2226 | 1113 |
| Water | 66 | 151 | 98 | 171 |
| **Ligands** |  |  |  |  |
| BMA | 2 | 1 | 2 | 1 |
| NAG | 40 | 20 | 40 | 20 |
| FUC | 2 | 1 | 2 | 1 |
| ZN | 2 | 1 | 2 | 1 |
| CC (ligands) | 0.67 | 0.63 | 0.68 | 0.66 |
| **B-factor (min/max/mean)** |  |  |  |  |
| Protein | 49.69/203.09/91.37 | 43.05/161.55/79.34 | 41.32/184.67/81.06 | 40.42/146.83/68.48 |
| Ligand | 84.58/197.39/124.52 | 66.11/151.60/106.69 | 74.15/180.56/11.94 | 63.35/140.87/97.33 |
| Water | 58.28/86.38/74.02 | 56.53/85.99/70.96 | 51.78/80.45/66.94 | 52.20/87.41/65.17 |
| **Model ID** |  |  |  |  |
| EMDB ID | EMD-70928 | EMD-70927 | EMD-70930 | EMD-70929 |
| PDB ID | 9OW4 | 9OW3 | 9OW6 | 9OW5 |

**Supplemental Figure 1.**
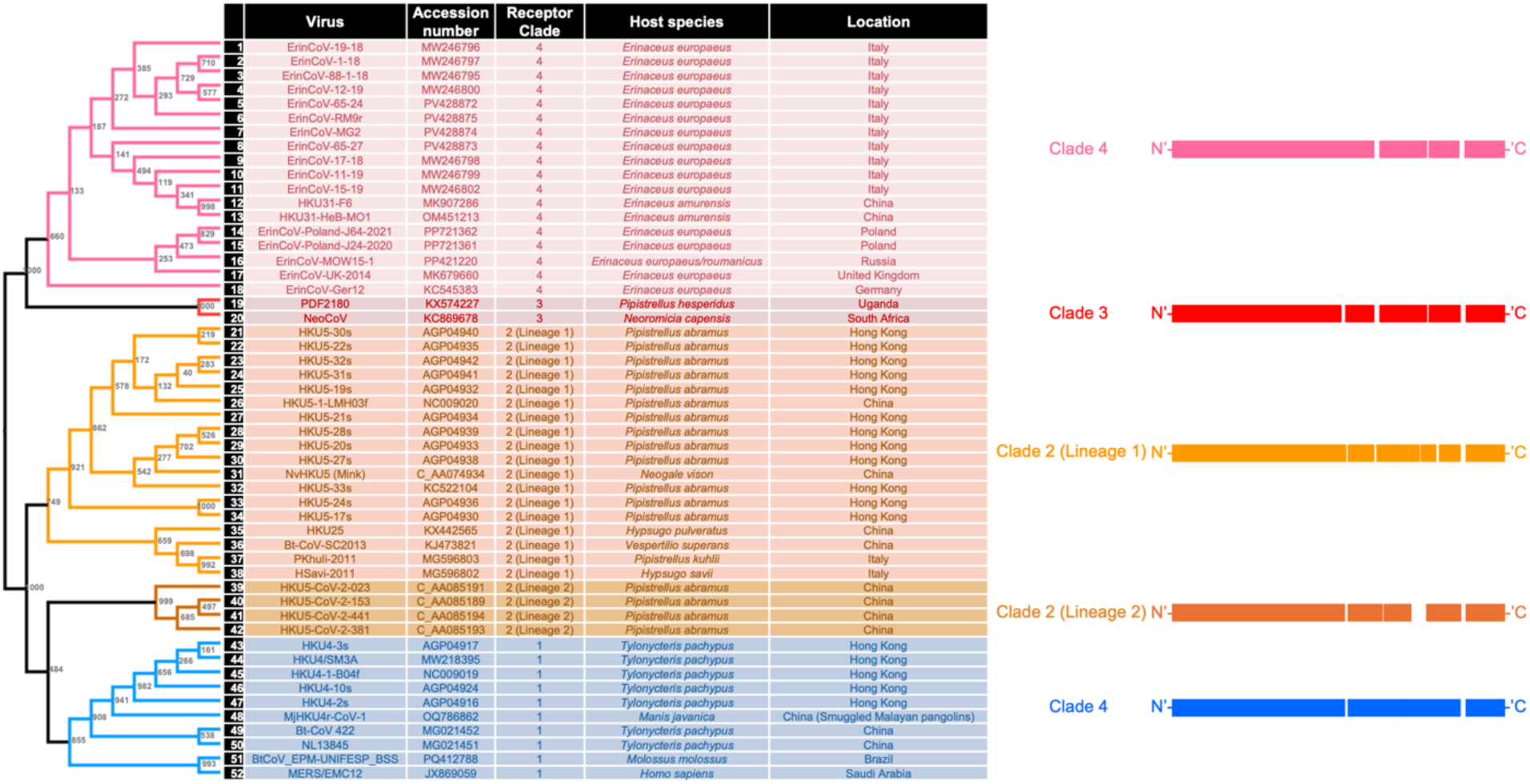
Phylogenetic similarity and schematic overview of merbecovirus RBDs and clades. All viruses in this study are listed in the table.

**Supplemental Figure 2.**
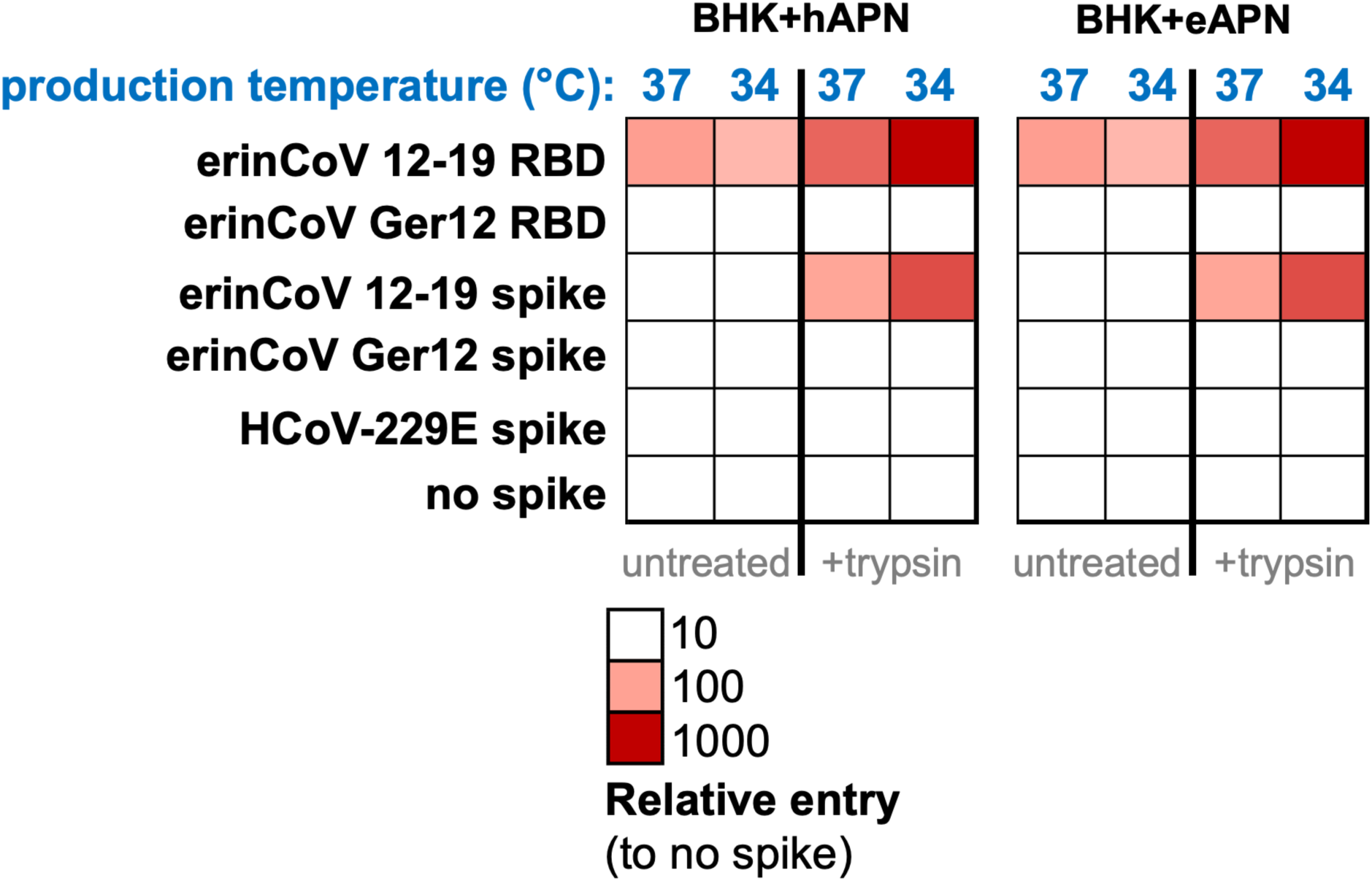
Viral pseudotypes are more infectious when produced at lower temperatures. During pseudotype production, 293T cells were moved to 34°C incubators following spike transfection and maintained at this temperature until pseudotype collection. Resulting pseudotypes from batches grown at 34 or 37°C were used to infect BHK cells expressing human or hedgehog APN.

**Supplemental Figure 3:**
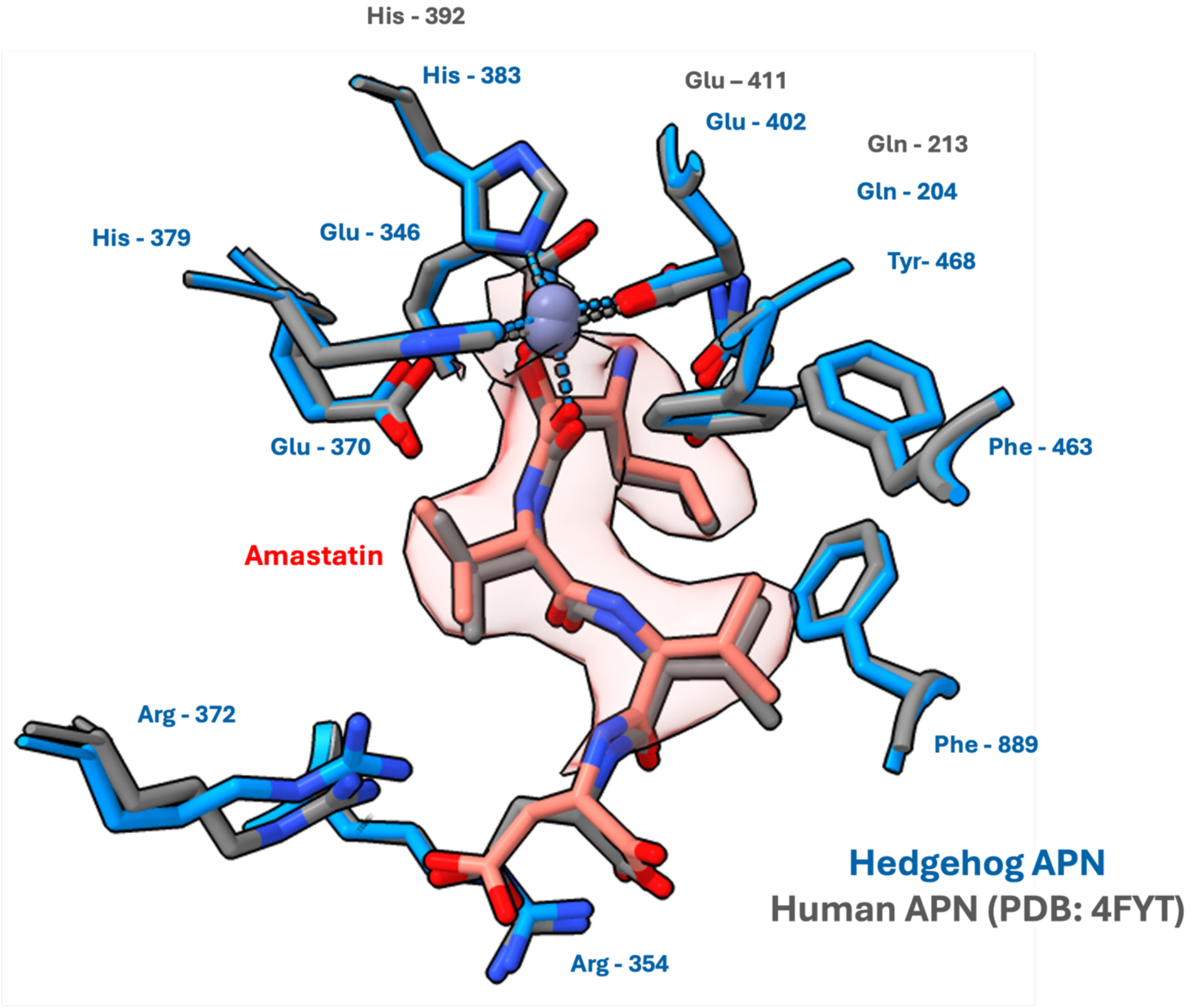
The amastatin binding site in hedgehog and human APN. Atomic structure of amastatin binding site. Residues and amastatin from the human APN-amastatin complex (PDB: 4FYT) are colored grey. In the hedgehog APN-amastatin complex (hedgehog APN – ErinCoV-Ger12 complex), residues from the hedgehog APN are colored blue and amastatin is colored salmon. Zinc ions are displayed as grey spheres. Only density around the bound amastatin from the locally refined cryo-EM map of the hedgehog – ErinCoV-Ger12 complex is shown.

**Supplemental Figure 4:**
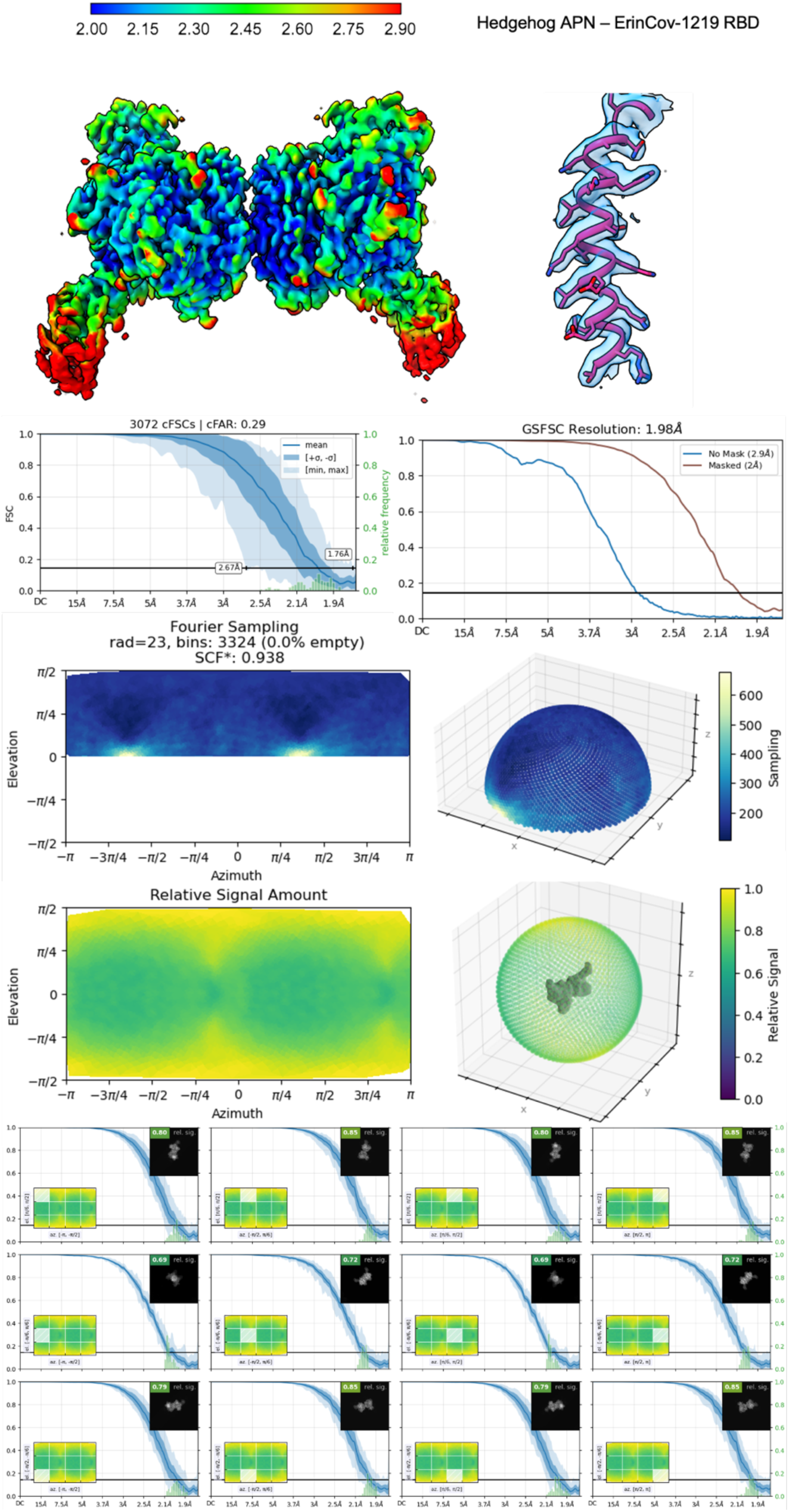
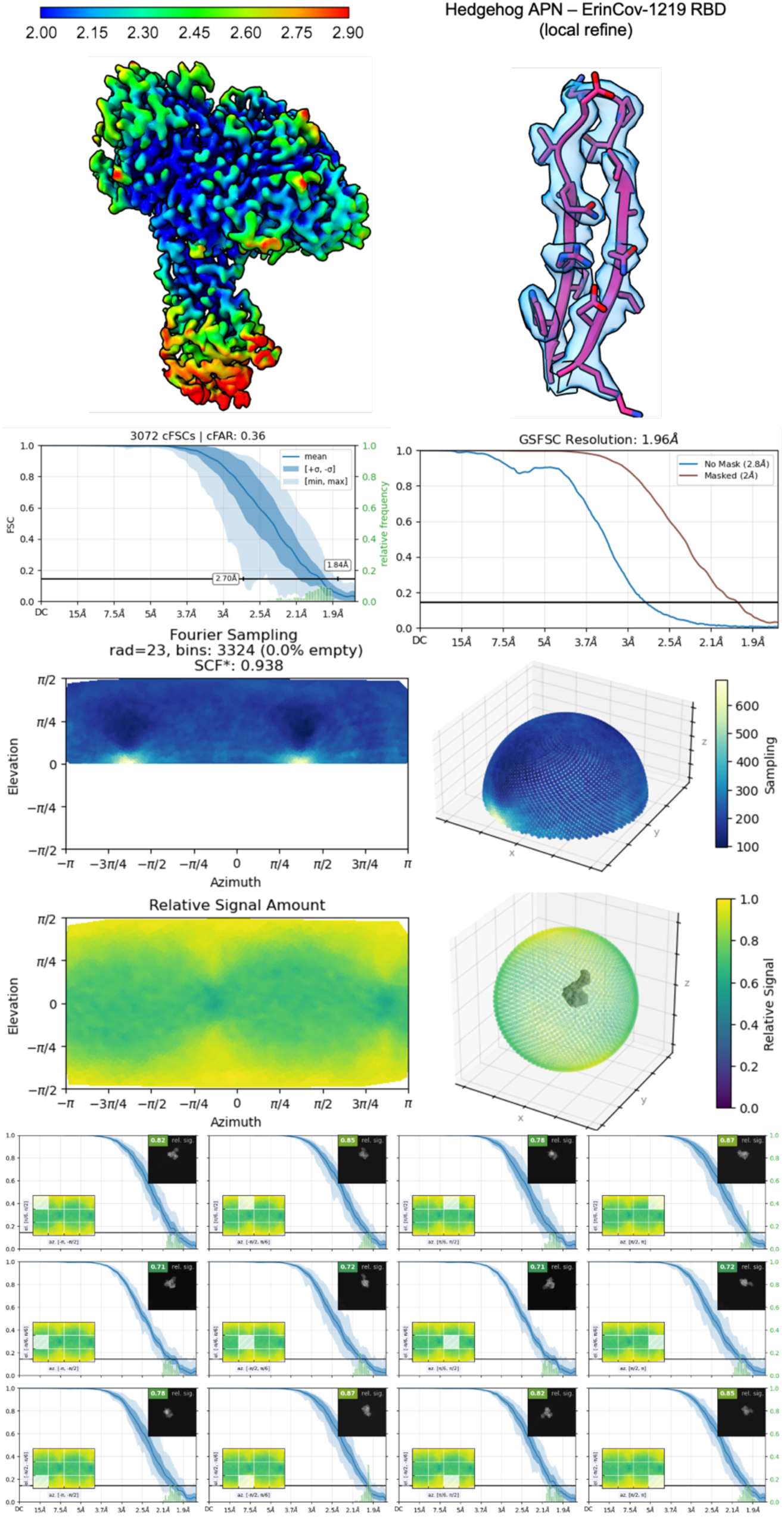
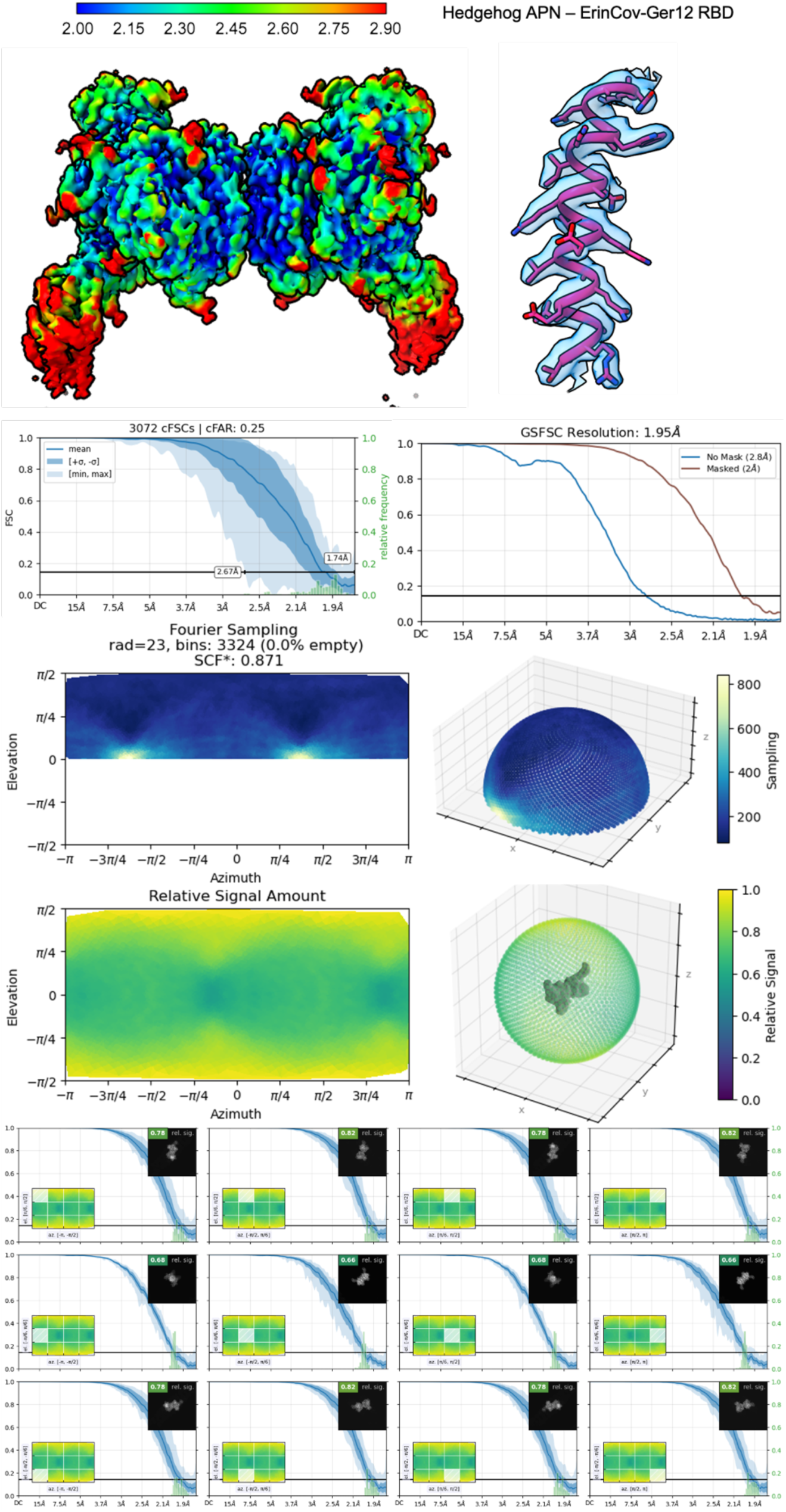
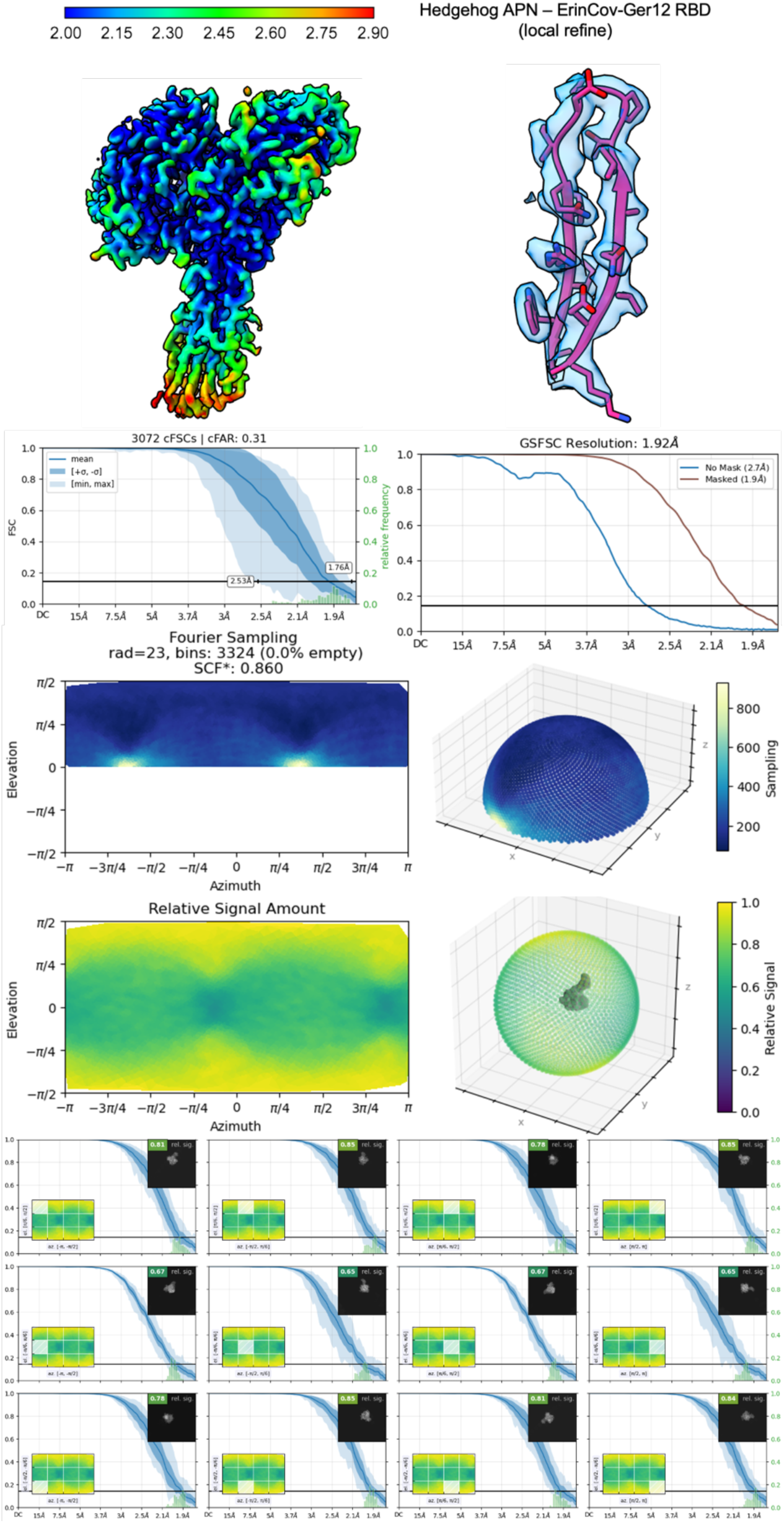
Orientational diagnostics of all the deposited cryo-EM structures.

**Supplemental Figure 5.**
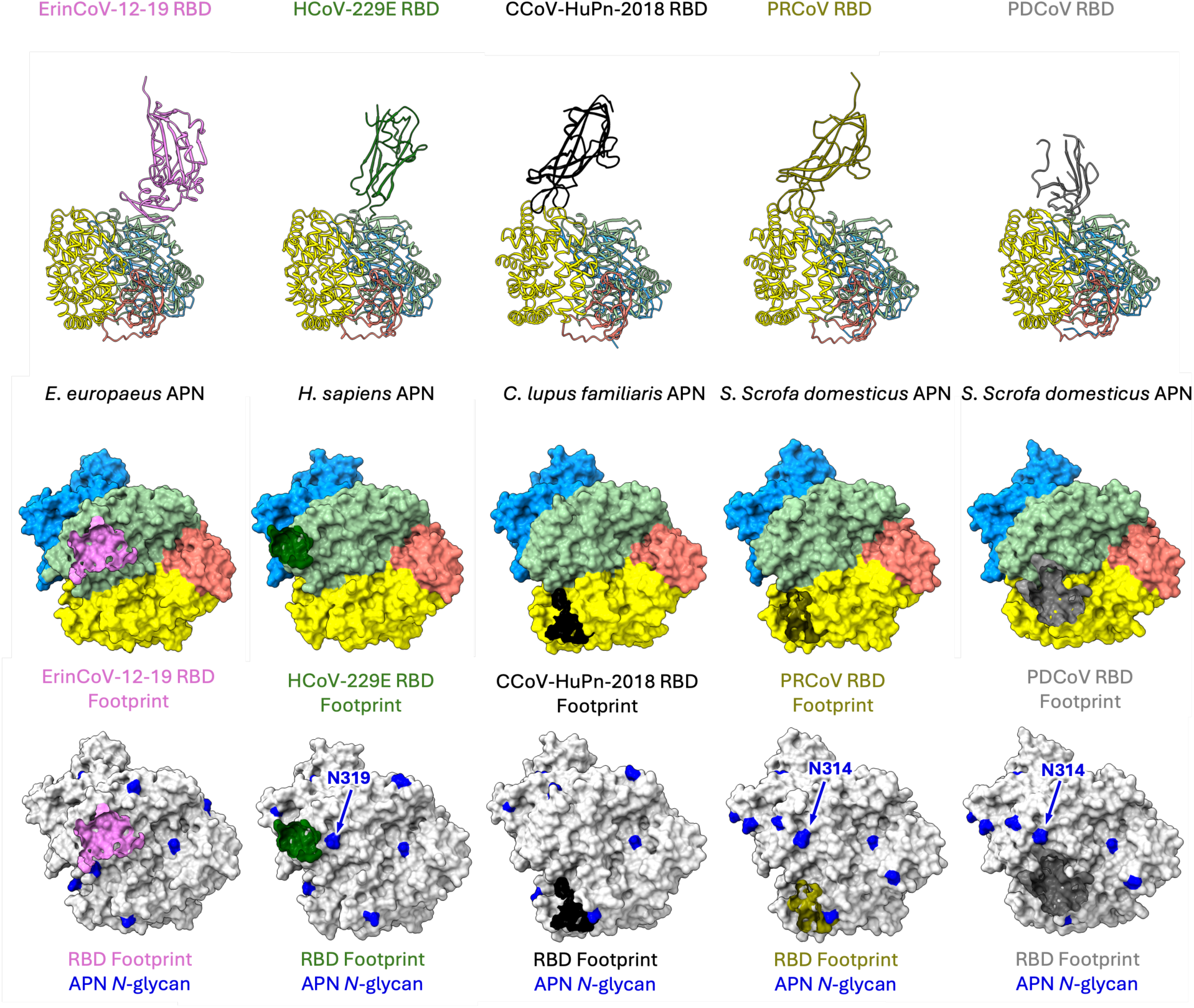
RBD interactions for coronaviruses that use APN. RBD:APN structures and RBD binding footprints for each of the following complexes are shown: **(i)** ErinCoV-12-19:eAPN (PDB:9OW3), **(ii)** HCoV-229E:hAPN (PDB: 6ATK), **(iii)** CCoV-HuPn-2018:cAPN (PDB: 7U0L), **(iv)** PRCoV:pAPN (PDB:4F5C) and **(v)** PDCoV:pAPN (PDB: 7VPP). In the top two panels, the APN structures and surfaces are colored by domain (Domain I: blue, Domain II: green, Domain III: salmon, Domain IV: yellow). In the bottom panel, the APN surface is colored grey and the *N-*glycosylated asparagines are colored blue. In the middle and bottom panels, the RBD footprint is overlayed on the APN surface and colored as in the top panel. The *N-*glycans found at the equivalent of K310 in hedgehog APN (Zone 3) are labeled.

**Supplemental Figure 6:**
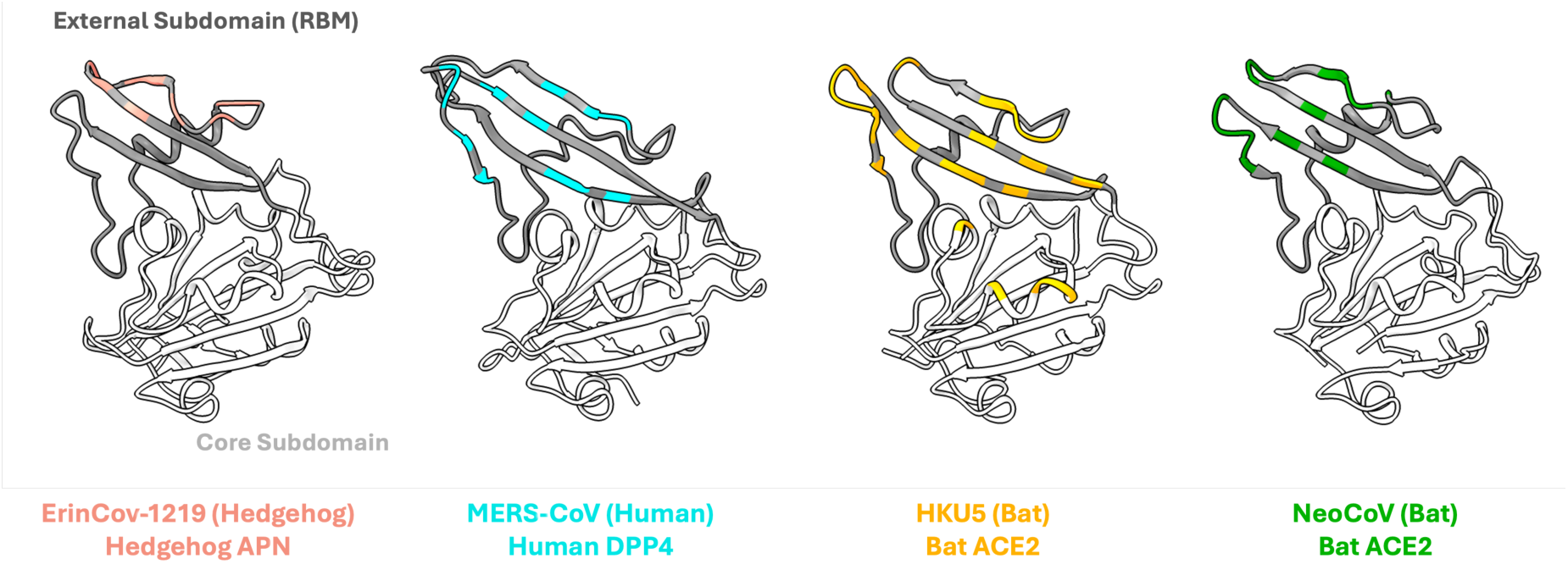
Merbecovirus receptor binding domains. The receptor binding domains only from each of the following merbecovirus RBD–receptor complexes are shown: **(i)** ErinCoV-12-19-eAPN (this work, PDB: 9OW3), **(ii)** MERS-CoV-hDPP4 (PDB: 4KR0), **(iii)** HKU5-bACE2 (PDB: 9D32) and **(iv)** NeoCoV-bACE2 (PDB: 7WPO). The receptor binding subdomain/receptor binding motif is colored dark grey, and the core subdomain is colored light grey. RBD residues that are in contact with the bound receptor are colored orange, cyan, yellow and green.

**Supplemental Figure 7:**
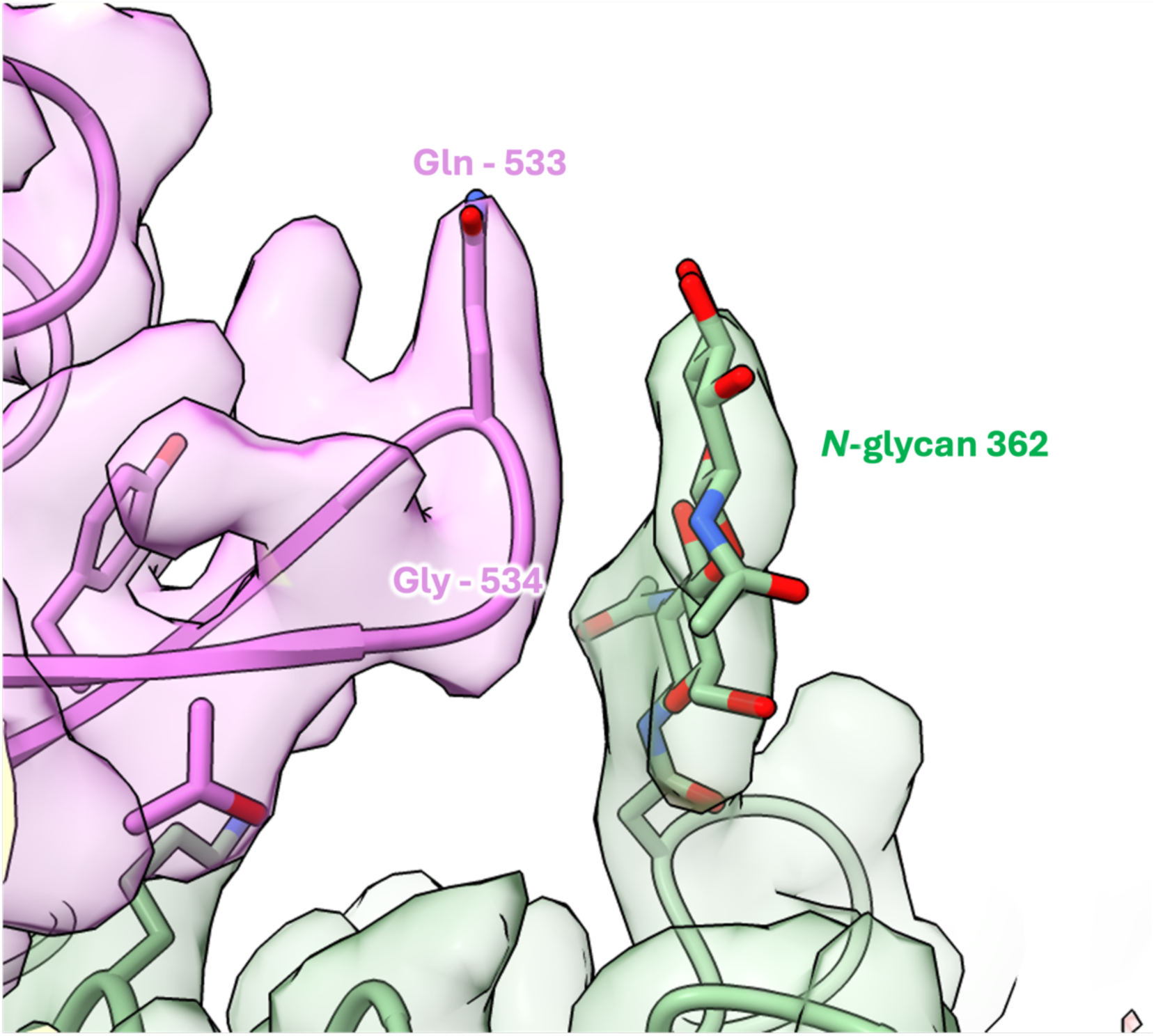
*N-*glycan at the ErinCoV-RBD – hedgehog APN interface. Atomic model of the ErinCoV-12-19 RBD – hedgehog APN complex.

**Supplemental Figure 8.**
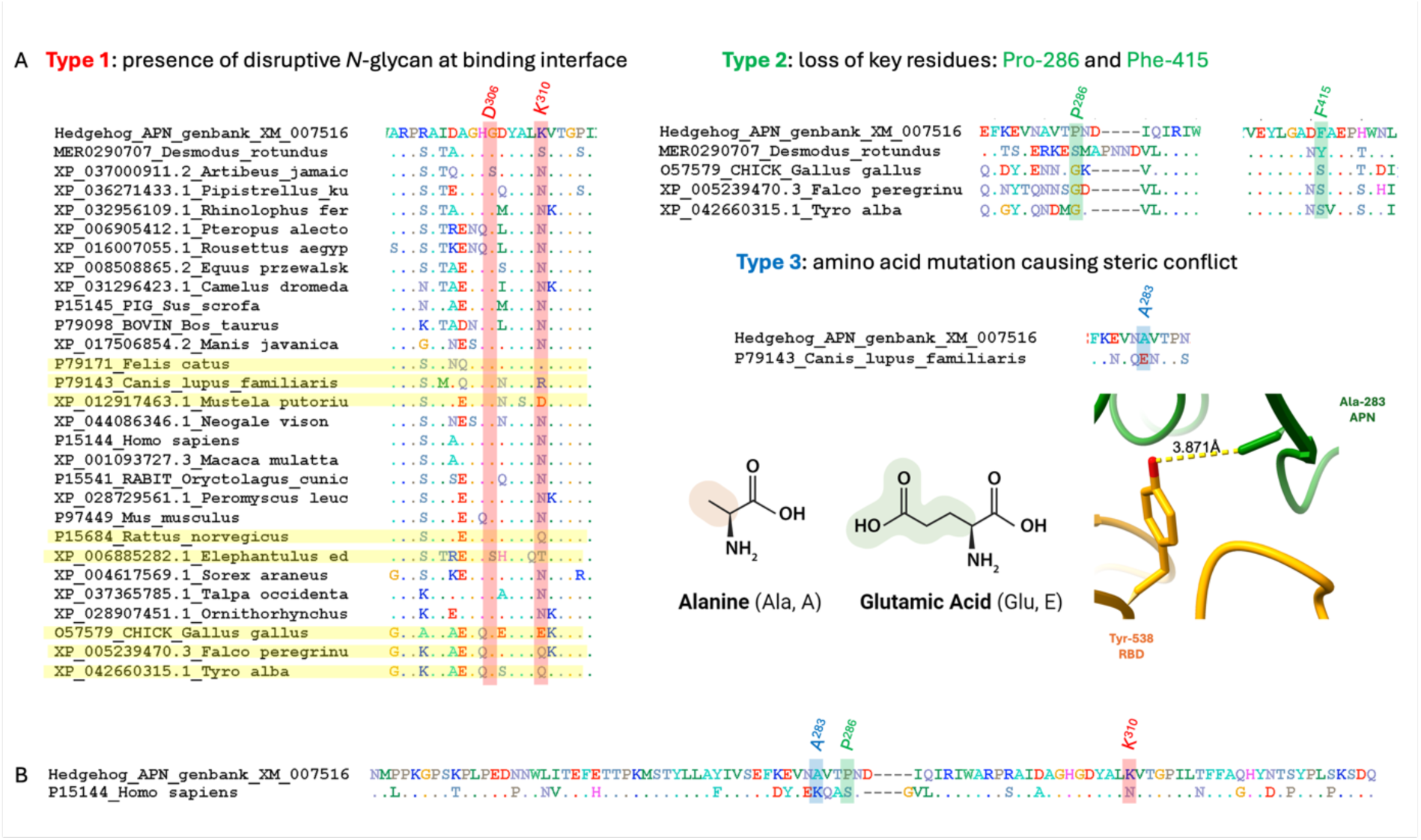
Variability among the tested APN sequences with the potential to block ErinCoV RBD binding. **(A)** There are three types of structural features that influence ErinCoVs’ ability to recognize APNs from other species. **(B)** Human APNs have characteristics from all three groups.

**Supplemental Figure 9:**
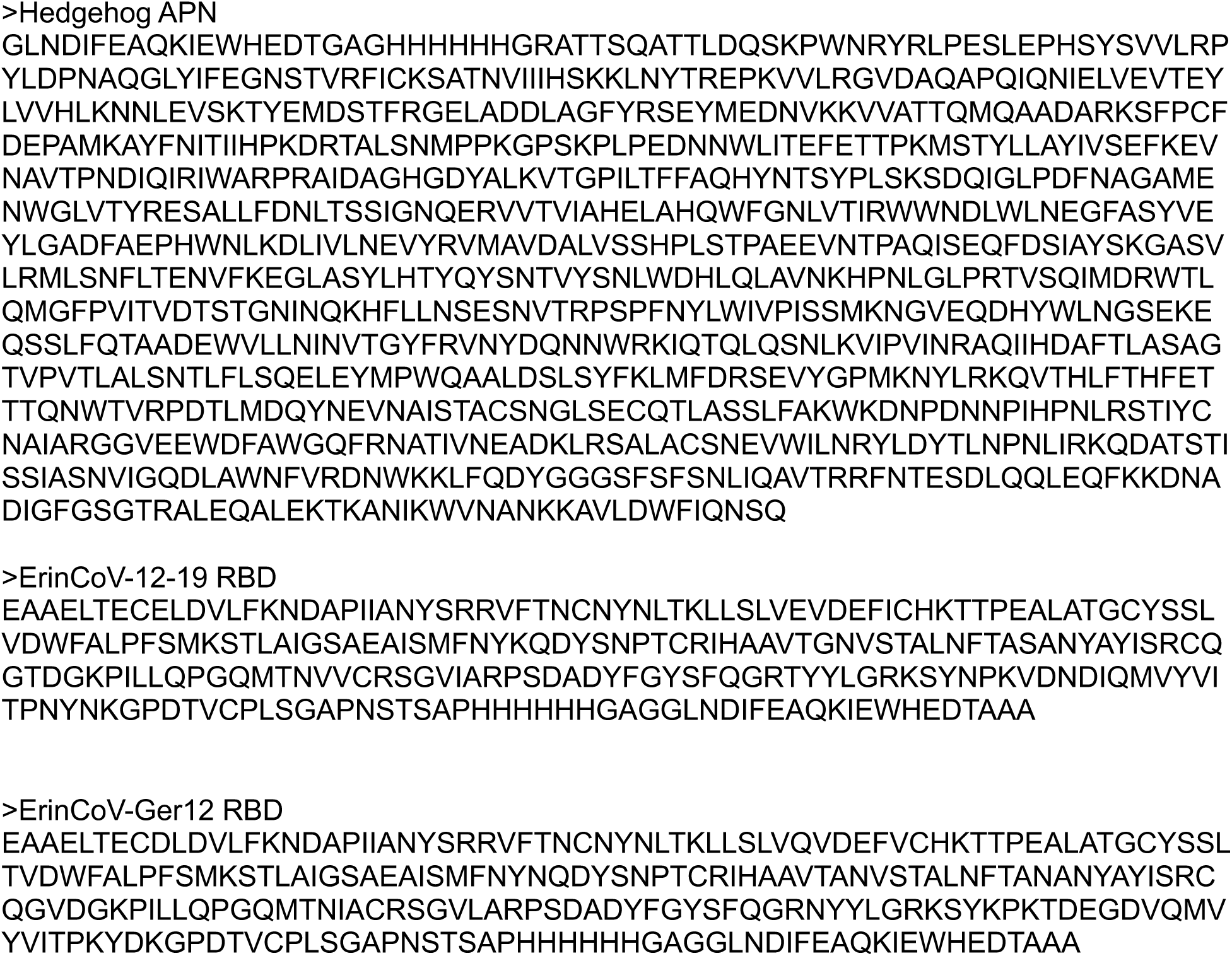
Amino acid sequences of all the constructs used in the cryo-EM and BLI binding studies.

**Supplemental Figure 10:**
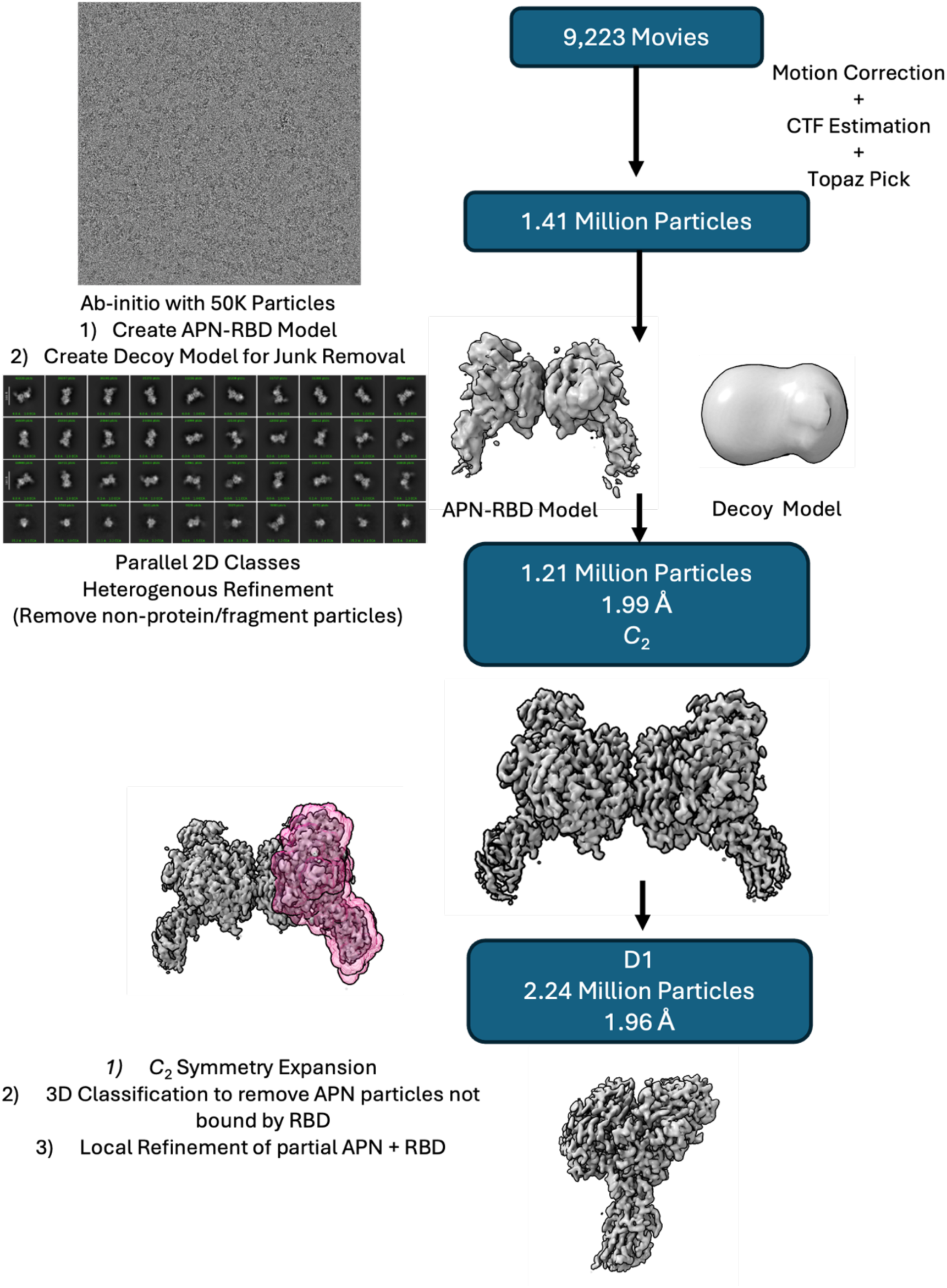
Cryo-EM flowchart of the hedgehog APN – ErinCoV-12-19 complex.

**Supplemental Figure 11:**
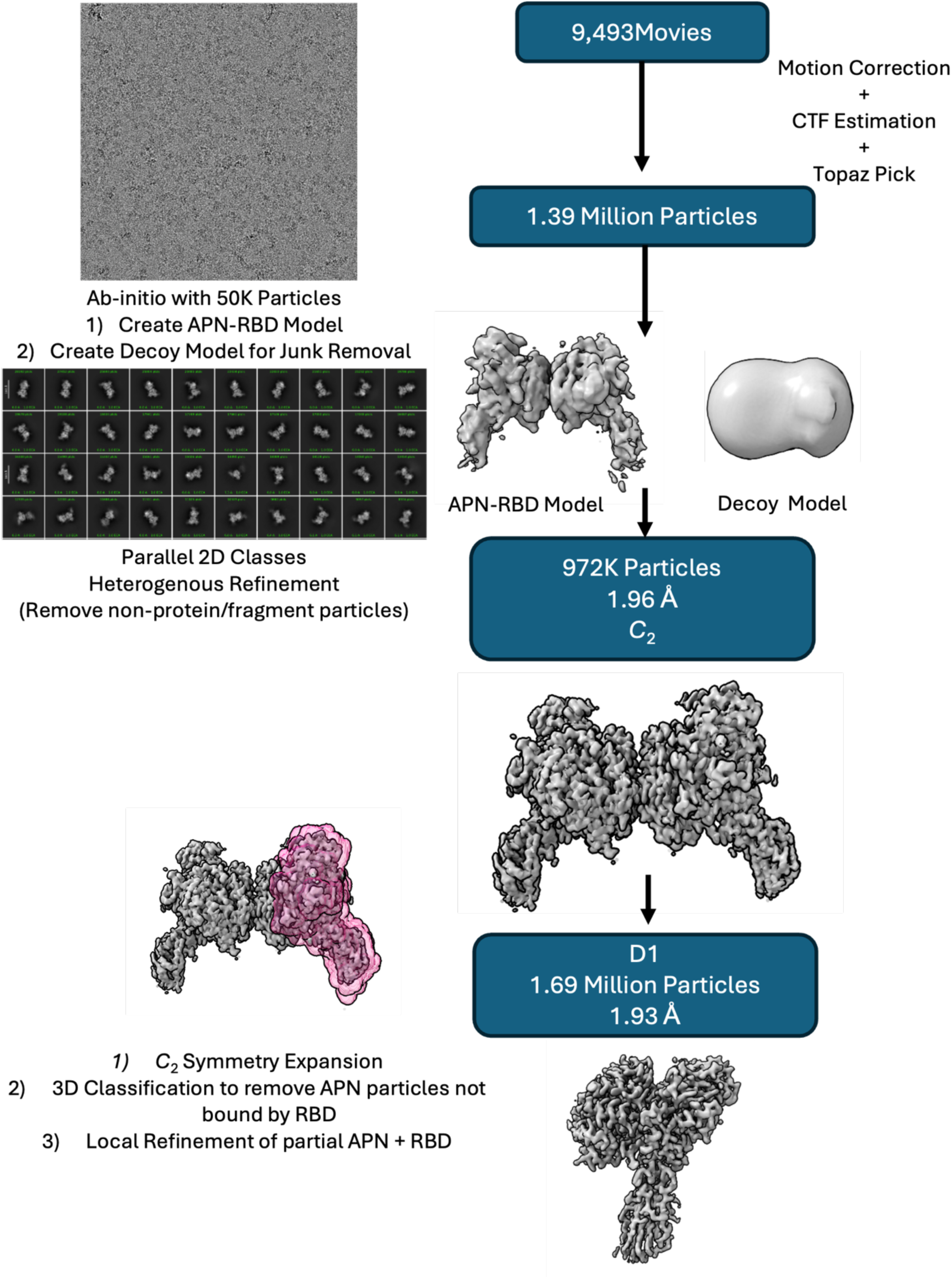
Cryo-EM flowchart of the hedgehog APN – ErinCoV-Ger12 complex.

**Supplemental Figure 12.**
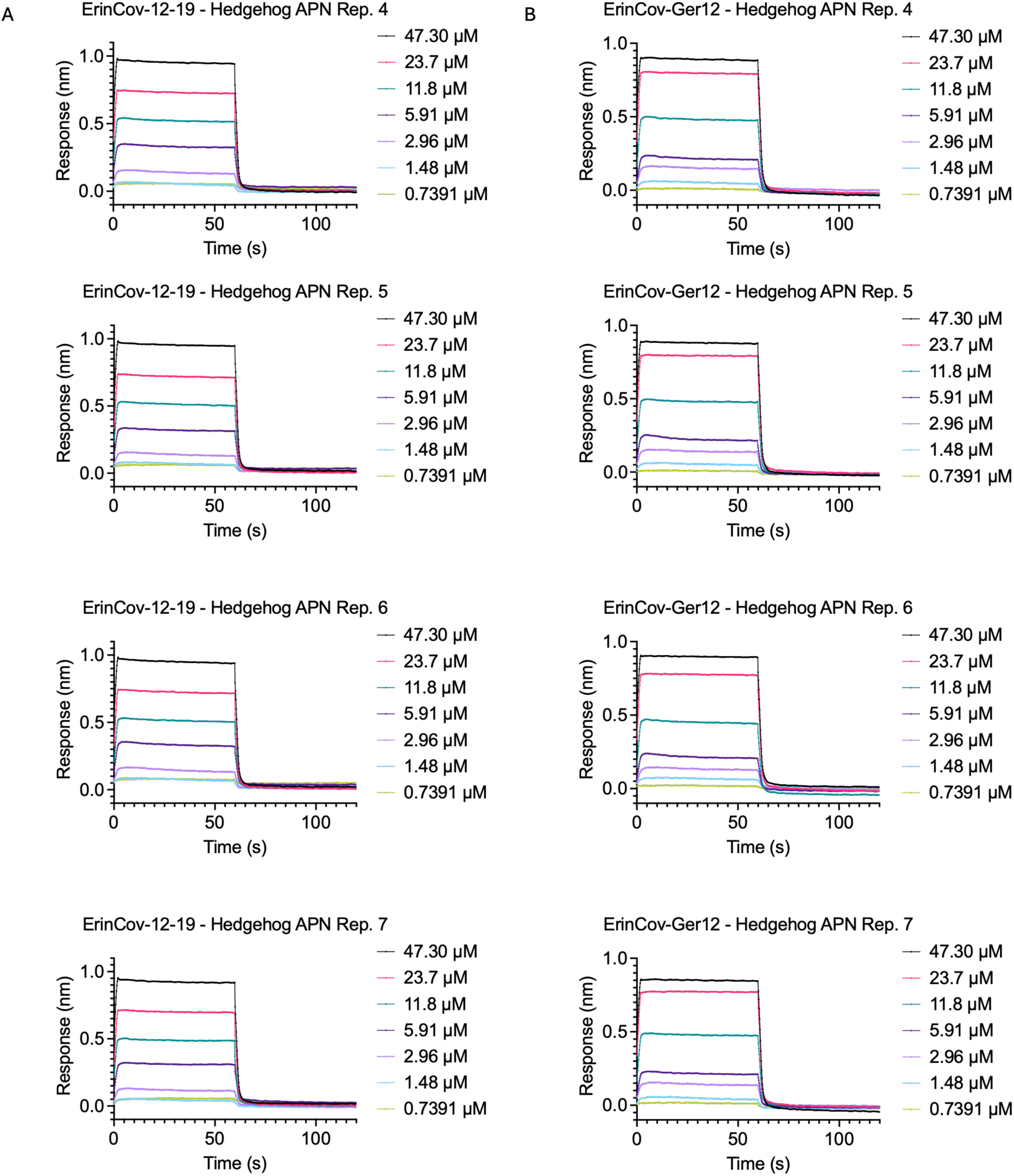
ErinCoV RBD – hedgehog APN binding sensorgrams. **(A)** BLI binding sensorgrams with the ErinCoV-12-19 RBD on the biosensor and hedgehog APN in solution **(B)** BLI binding sensorgrams with the ErinCoV-Ger12 RBD on the biosensor and hedgehog APN in solution

## ACKNOWLEDGEMENTS

We would like to thank Dr. Alex Cohen for providing comments and insights on the work. Research reported in this publication was supported by the National Institute of Allergy and Infectious Diseases of the National Institutes of Health (NIAID/NIH) under Award Numbers R01AI179720 to ML, R01 AI167966 and CEIRS Contract 75N93021C00014 to RSB, and T32AI007025 to VAJ. TNS is a Searle Scholars Fellow. This work was supported in part by the intramural program of the National Institute of Allergy and Infectious Diseases. The content is solely the responsibility of the author and does not necessarily represent the official views of the National Institutes of Health.

## AUTHOR CONTRIBUTIONS

MJ, VAJ, ZZ, NJC, LDS, ELK, BLM, AC, IDB, AM, LDS, SNS, AC, TNS, and ML performed experiments and analysis. MJ, VAJ, ZZ, NJC, and ML generated figures. AM, GV, BAH, CBW, RSB, JMR, and ML provided resources, supervision, and funding. ML developed the project concept and coordinated collaboration. All authors contributed to the manuscript and approved the final version of the text and figures.

## REFERENCES

1. Zaki, A. M., Van Boheemen, S., Bestebroer, T. M., Osterhaus, A. D. M. E. & Fouchier, R. A. M. Isolation of a Novel Coronavirus from a Man with Pneumonia in Saudi Arabia. N. Engl. J. Med. 367, 1814–1820 (2012).

2. World Health Organization. Middle East respiratory syndrome coronavirus - Kingdom of Saudi Arabia; Disease Outbreak News. https://www.who.int/emergencies/disease-outbreak-news/item/2025-DON569 (2025).

3. Azhar, E. I. et al. Evidence for Camel-to-Human Transmission of MERS Coronavirus. N. Engl. J. Med. 370, 2499–2505 (2014).

4. Woo, P. C., Lau, S. K., Li, K. S., Tsang, A. K. & Yuen, K.-Y. Genetic relatedness of the novel human group C betacoronavirus to Tylonycteris bat coronavirus HKU4 and Pipistrellus bat coronavirus HKU5. Emerg. Microbes Infect. 1, e35 (2012).

5. Ithete, N. L. et al. Close Relative of Human Middle East Respiratory Syndrome Coronavirus in Bat, South Africa. Emerg. Infect. Dis. 19, 1697–1699 (2013).

6. Wu, Z. et al. ORF8-Related Genetic Evidence for Chinese Horseshoe Bats as the Source of Human Severe Acute Respiratory Syndrome Coronavirus. J. Infect. Dis. 213, 579–583 (2016).

7. Anthony, S. J. et al. Further Evidence for Bats as the Evolutionary Source of Middle East Respiratory Syndrome Coronavirus. mBio 8, e00373–17 (2017).

8. Moreno, A. et al. Detection and full genome characterization of two beta CoV viruses related to Middle East respiratory syndrome from bats in Italy. Virol. J. 14, 239 (2017).

9. Lau, S. K. P. et al. Receptor Usage of a Novel Bat Lineage C Betacoronavirus Reveals Evolution of Middle East Respiratory Syndrome-Related Coronavirus Spike Proteins for Human Dipeptidyl Peptidase 4 Binding. J. Infect. Dis. 218, 197–207 (2018).

10. Speranskaya, A. S. et al. Identification and Genetic Characterization of MERS-Related Coronavirus Isolated from Nathusius’ Pipistrelle (Pipistrellus nathusii) near Zvenigorod (Moscow Region, Russia). Int. J. Environ. Res. Public. Health 20, 3702 (2023).

11. Silvério, B. S. et al. Coronavirus Cryptic Landscape and Draft Genome of a Novel CoV Clade Related to MERS From Bats Circulating in Northeastern Brazil. J. Med. Virol. 97, e70173 (2025).

12. Chen, J. et al. A bat MERS-like coronavirus circulates in pangolins and utilizes human DPP4 and host proteases for cell entry. Cell 186, 850–863.e16 (2023).

13. Zhao, J. et al. Farmed fur animals harbour viruses with zoonotic spillover potential. Nature 634, 228–233 (2024).

14. Corman, V. M. et al. Characterization of a Novel Betacoronavirus Related to Middle East Respiratory Syndrome Coronavirus in European Hedgehogs. J. Virol. 88, 717–724 (2014).

15. Bininda-Emonds, O. R. P. et al. The delayed rise of present-day mammals. Nature 446, 507–512 (2007).

16. Woo, P. C. Y., Lau, S. K. P., Huang, Y. & Yuen, K.-Y. Coronavirus diversity, phylogeny and interspecies jumping. Exp. Biol. Med. Maywood NJ 234, 1117–1127 (2009).

17. Tsagkogeorga, G., Parker, J., Stupka, E., Cotton, J. A. & Rossiter, S. J. Phylogenomic Analyses Elucidate the Evolutionary Relationships of Bats. Curr. Biol. 23, 2262–2267 (2013).

18. Monchatre-Leroy, E. et al. Identification of Alpha and Beta Coronavirus in Wildlife Species in France: Bats, Rodents, Rabbits, and Hedgehogs. Viruses 9, 364 (2017).

19. Beissat, K. et al. Coding-complete genome sequences of hedgehog coronavirus isolated from Erinaceus europaeus in France. Microbiol. Resour. Announc. 0, e00157–25 (2025).

20. Saldanha, I. F. et al. Extension of the known distribution of a novel clade C betacoronavirus in a wildlife host. Epidemiol. Infect. 147, e169 (2019).

21. Lau, S. K. P. et al. Identification of a Novel Betacoronavirus (Merbecovirus) in Amur Hedgehogs from China. Viruses 11, 980 (2019).

22. He, W.-T. et al. Virome characterization of game animals in China reveals a spectrum of emerging pathogens. Cell 185, 1117–1129.e8 (2022).

23. Delogu, M. et al. Eco-Virological Preliminary Study of Potentially Emerging Pathogens in Hedgehogs (Erinaceus europaeus) Recovered at a Wildlife Treatment and Rehabilitation Center in Northern Italy. Animals 10, 407 (2020).

24. De Sabato, L. et al. Erinaceus coronavirus persistence in hedgehogs (Erinaceus europaeus) in a non-invasive, in vivo, experimental setting. Front. Vet. Sci. 10, 1213990 (2023).

25. Pomorska-Mól, M., Ruszkowski, J. J., Gogulski, M. & Domanska-Blicharz, K. First detection of Hedgehog coronavirus 1 in Poland. Sci. Rep. 12, 2386 (2022).

26. Lukina-Gronskaya, A. V. et al. Novel coronaviruses and mammarenaviruses of hedgehogs from Russia including the comparison of viral communities of hibernating and active specimens. Front. Vet. Sci. 11, 1486635 (2024).

27. Cruz, A. V. S. et al. Genomic characterization and cross-species transmission potential of hedgehog coronavirus. One Health 19, 100940 (2024).

28. Ruszkowski, J. J., Hetman, M., Turlewicz-Podbielska, H. & Pomorska-Mól, M. Hedgehogs as a Potential Source of Zoonotic Pathogens—A Review and an Update of Knowledge. Anim. Open Access J. MDPI 11, 1754 (2021).

29. Hubert, P., Julliard, R., Biagianti, S. & Poulle, M.-L. Ecological factors driving the higher hedgehog (*Erinaceus europeaus*) density in an urban area compared to the adjacent rural area. Landsc. Urban Plan. 103, 34–43 (2011).

30. van de Poel, J. L., Dekker, J. & van Langevelde, F. Dutch hedgehogs Erinaceus europaeus are nowadays mainly found in urban areas, possibly due to the negative Effects of badgers Meles meles. Wildl. Biol. 21, wlb.00855 (2015).

31. Fan, C. et al. A Human DPP4-Knockin Mouse’s Susceptibility to Infection by Authentic and Pseudotyped MERS-CoV. Viruses 10, 448 (2018).

32. Tailor, N. et al. Generation and Characterization of a SARS-CoV-2-Susceptible Mouse Model Using Adeno-Associated Virus (AAV6.2FF)-Mediated Respiratory Delivery of the Human ACE2 Gene. Viruses 15, 85 (2022).

33. Chau, L. F. et al. Human ACE2 transgenic pigs are susceptible to SARS-CoV-2 and develop COVID-19-like disease. Nat. Commun. 16, 766 (2025).

34. Alfajaro, M. M. et al. HKU5 bat merbecoviruses use divergent mechanisms to engage bat and mink ACE2 as entry receptors. 2025.02.12.637862 Preprint at 10.1101/2025.02.12.637862 (2025).

35. V’kovski, P., Kratzel, A., Steiner, S., Stalder, H. & Thiel, V. Coronavirus biology and replication: implications for SARS-CoV-2. Nat. Rev. Microbiol. 19, 155–170 (2021).

36. Everest, H., Stevenson-Leggett, P., Bailey, D., Bickerton, E. & Keep, S. Known Cellular and Receptor Interactions of Animal and Human Coronaviruses: A Review. Viruses 14, 351 (2022).

37. Walls, A. C. et al. Tectonic conformational changes of a coronavirus spike glycoprotein promote membrane fusion. Proc. Natl. Acad. Sci. U. S. A. 114, 11157–11162 (2017).

38. Cai, Y. et al. Distinct conformational states of SARS-CoV-2 spike protein. Science 369, 1586–1592 (2020).

39. Jackson, C. B., Farzan, M., Chen, B. & Choe, H. Mechanisms of SARS-CoV-2 entry into cells. Nat. Rev. Mol. Cell Biol. 23, 3–20 (2022).

40. Li, W. et al. Angiotensin-converting enzyme 2 is a functional receptor for the SARS coronavirus. Nature 426, 450–454 (2003).

41. Zhou, P. et al. A pneumonia outbreak associated with a new coronavirus of probable bat origin. Nature 579, 270–273 (2020).

42. Hofmann, H. et al. Human coronavirus NL63 employs the severe acute respiratory syndrome coronavirus receptor for cellular entry. Proc. Natl. Acad. Sci. U. S. A. 102, 7988– 7993 (2005).

43. Yeager, C. L. et al. Human aminopeptidase N is a receptor for human coronavirus 229E. Nature 357, 420–422 (1992).

44. Raj, V. S. et al. Dipeptidyl peptidase 4 is a functional receptor for the emerging human coronavirus-EMC. Nature 495, 251–254 (2013).

45. Yang, Y. et al. Receptor usage and cell entry of bat coronavirus HKU4 provide insight into bat-to-human transmission of MERS coronavirus. Proc. Natl. Acad. Sci. U. S. A. 111, 12516–12521 (2014).

46. Saunders, N. et al. TMPRSS2 is a functional receptor for human coronavirus HKU1. Nature 624, 207–214 (2023).

47. Dveksler, G. S. et al. Cloning of the mouse hepatitis virus (MHV) receptor: expression in human and hamster cell lines confers susceptibility to MHV. J. Virol. 65, 6881–6891 (1991).

48. Dufloo, J. et al. Dipeptidase 1 is a functional receptor for coronavirus PHEV. 2025.01.09.632101 Preprint at 10.1101/2025.01.09.632101 (2025).

49. Lang, Y. et al. Coronavirus hemagglutinin-esterase and spike proteins coevolve for functional balance and optimal virion avidity. Proc. Natl. Acad. Sci. U. S. A. 117, 25759–25770 (2020).

50. Yokomori, K., Banner, L. R. & Lai, M. M. C. Heterogeneity of gene expression of the hemagglutinin-esterase (HE) protein of murine coronaviruses. Virology 183, 647–657 (1991).

51. de Groot, R. J. Structure, function and evolution of the hemagglutinin-esterase proteins of corona- and toroviruses. Glycoconj. J. 23, 59–72 (2006).

52. Catanzaro, N. J. et al. ACE2 from Pipistrellus abramus bats is a receptor for HKU5 coronaviruses. Nat. Commun. 16, 4932 (2025).

53. Lau, S. K. P. et al. Isolation of MERS-related coronavirus from lesser bamboo bats that uses DPP4 and infects human-DPP4-transgenic mice. Nat. Commun. 12, 216 (2021).

54. Wang, Q. et al. Bat origins of MERS-CoV supported by bat coronavirus HKU4 usage of human receptor CD26. Cell Host Microbe 16, 328–337 (2014).

55. Xia, L.-Y. et al. Isolation and characterization of a pangolin-borne HKU4-related coronavirus that potentially infects human-DPP4-transgenic mice. Nat. Commun. 15, 1048 (2024).

56. Xiong, Q. et al. Close relatives of MERS-CoV in bats use ACE2 as their functional receptors. Nature 612, 748–757 (2022).

57. Ji, W. et al. Structures of a deltacoronavirus spike protein bound to porcine and human receptors. Nat. Commun. 13, 1467 (2022).

58. Wong, A. H. M. et al. Receptor-binding loops in alphacoronavirus adaptation and evolution. Nat. Commun. 8, 1735 (2017).

59. Tortorici, M. A. et al. Structure, receptor recognition, and antigenicity of the human coronavirus CCoV-HuPn-2018 spike glycoprotein. Cell 185, 2279–2291.e17 (2022).

60. Reguera, J. et al. Structural bases of coronavirus attachment to host aminopeptidase N and its inhibition by neutralizing antibodies. PLoS Pathog. 8, e1002859 (2012).

61. St-Jean, J. R. et al. Recovery of a Neurovirulent Human Coronavirus OC43 from an Infectious cDNA Clone. J. Virol. 80, 3670–3674 (2006).

62. Fowler, P. A. & Racey, P. A. Daily and seasonal cycles of body temperature and aspects of heterothermy in the hedgehog Eriuaceus europaeus. J. Comp. Physiol. [B] 160, 299–307 (1990).

63. Laporte, M. et al. The SARS-CoV-2 and other human coronavirus spike proteins are fine-tuned towards temperature and proteases of the human airways. PLoS Pathog. 17, e1009500 (2021).

64. Letko, M., Marzi, A. & Munster, V. Functional assessment of cell entry and receptor usage for SARS-CoV-2 and other lineage B betacoronaviruses. Nat. Microbiol. 5, 562–569 (2020).

65. Guo, H. et al. ACE2-Independent Bat Sarbecovirus Entry and Replication in Human and Bat Cells. mBio 13, e02566–22 (2022).

66. Guo, H. et al. Isolation of ACE2-dependent and -independent sarbecoviruses from Chinese horseshoe bats. J. Virol. 97, e00395–23 (2023).

67. Seifert, S. N. et al. An ACE2-dependent Sarbecovirus in Russian bats is resistant to SARS-CoV-2 vaccines. PLOS Pathog. 18, e1010828 (2022).

68. Tse, A. L. et al. Distinct pathways for evolution of enhanced receptor binding and cell entry in SARS-like bat coronaviruses. PLOS Pathog. 20, e1012704 (2024).

69. Jin, M. et al. Human coronavirus HKU1 spike structures reveal the basis for sialoglycan specificity and carbohydrate-promoted conformational changes. Nat. Commun. 16, 4158 (2025).

70. Beier, K. T. et al. Anterograde or retrograde transsynaptic labeling of CNS neurons with vesicular stomatitis virus vectors. Proc. Natl. Acad. Sci. U. S. A. 108, 15414–15419 (2011).

71. Plavec, Z. et al. SARS-CoV-2 Production, Purification Methods and UV Inactivation for Proteomics and Structural Studies. Viruses 14, 1989 (2022).

72. Wong, A. H. M., Zhou, D. & Rini, J. M. The X-ray crystal structure of human aminopeptidase N reveals a novel dimer and the basis for peptide processing. J. Biol. Chem. 287, 36804– 36813 (2012).

73. Park, Y.-J. et al. Molecular basis of convergent evolution of ACE2 receptor utilization among HKU5 coronaviruses. Cell 188, 1711–1728.e21 (2025).

74. Lu, G. et al. Molecular basis of binding between novel human coronavirus MERS-CoV and its receptor CD26. Nature 500, 227–231 (2013).

75. Delmas, B. et al. Aminopeptidase N is a major receptor for the entero-pathogenic coronavirus TGEV. Nature 357, 417–420 (1992).

76. Delmas, B., Gelfi, J., Sjöström, H., Noren, O. & Laude, H. Further characterization of aminopeptidase-N as a receptor for coronaviruses. Adv. Exp. Med. Biol. 342, 293–298 (1993).

77. Tresnan, D. B., Levis, R. & Holmes, K. V. Feline aminopeptidase N serves as a receptor for feline, canine, porcine, and human coronaviruses in serogroup I. J. Virol. 70, 8669–8674 (1996).

78. Chen, J. et al. Bat-infecting merbecovirus HKU5-CoV lineage 2 can use human ACE2 as a cell entry receptor. Cell 188, 1729–1742.e16 (2025).

79. Wang, N. et al. A MERS-CoV-like mink coronavirus uses ACE2 as an entry receptor. Nature 642, 739–746 (2025).

80. Marr, C. R., Benlekbir, S. & Rubinstein, J. L. Fabrication of carbon films with ∼ 500nm holes for cryo-EM with a direct detector device. J. Struct. Biol. 185, 42–47 (2014).

81. Punjani, A., Rubinstein, J. L., Fleet, D. J. & Brubaker, M. A. cryoSPARC: algorithms for rapid unsupervised cryo-EM structure determination. Nat. Methods 14, 290–296 (2017).

82. Meng, E. C. et al. UCSF ChimeraX: Tools for structure building and analysis. Protein Sci. Publ. Protein Soc. 32, e4792 (2023).

83. Emsley, P. & Cowtan, K. Coot: model-building tools for molecular graphics. Acta Crystallogr. D Biol. Crystallogr. 60, 2126–2132 (2004).

84. Abramson, J. et al. Accurate structure prediction of biomolecular interactions with AlphaFold 3. Nature 630, 493–500 (2024).

85. Croll, T. I. ISOLDE: a physically realistic environment for model building into low-resolution electron-density maps. Acta Crystallogr. Sect. Struct. Biol. 74, 519–530 (2018).

86. Liebschner, D. et al. Macromolecular structure determination using X-rays, neutrons and electrons: recent developments in Phenix. Acta Crystallogr. Sect. Struct. Biol. 75, 861–877 (2019).

87. Pettersen, E. F. et al. UCSF ChimeraX: Structure visualization for researchers, educators, and developers. Protein Sci. Publ. Protein Soc. 30, 70–82 (2021).

88. Letko, M., Booiman, T., Kootstra, N., Simon, V. & Ooms, M. Identification of the HIV-1 Vif and Human APOBEC3G Protein Interface. Cell Rep. 13, 1789–1799 (2015).

89. Letko, M. et al. Adaptive Evolution of MERS-CoV to Species Variation in DPP4. Cell Rep. 24, 1730–1737 (2018).

90. Guindon, S. et al. New algorithms and methods to estimate maximum-likelihood phylogenies: assessing the performance of PhyML 3.0. Syst. Biol. 59, 307–321 (2010).

91. Lefort, V., Longueville, J.-E. & Gascuel, O. SMS: Smart Model Selection in PhyML. Mol. Biol. Evol. 34, 2422–2424 (2017).

92. Khaledian, E. et al. Sequence determinants of human-cell entry identified in ACE2-independent bat sarbecoviruses: A combined laboratory and computational network science approach. eBioMedicine 79, 103990 (2022).

93. Wei, J. et al. Genome-wide CRISPR Screens Reveal Host Factors Critical for SARS-CoV-2 Infection. Cell 184, 76–91.e13 (2021).

94. Whelan, S. P., Ball, L. A., Barr, J. N. & Wertz, G. T. Efficient recovery of infectious vesicular stomatitis virus entirely from cDNA clones. Proc. Natl. Acad. Sci. U. S. A. 92, 8388–8392 (1995).

